# Selective targeting of a histone-like silencer Sfx to the R6K conjugal transfer operon

**DOI:** 10.64898/2026.02.23.707533

**Authors:** Bing Wang, Ritika Gupta, Nathan Blaine, Barbare Khitiri, Catherine Jordan, Natalia Molotievskiy, David Dunlap, Laura Finzi, Irina Artsimovitch

## Abstract

Conjugative plasmids drive bacterial evolution and antibiotic resistance spread, yet their gene expression must be silenced to protect the host. A histone-like protein H-NS represses many mobile and sedentary xenogenes but fails to silence the conjugal transfer *vir* operon of R6K, a prototype IncX plasmid. Instead, R6K encodes its own H-NS homolog, Sfx, to repress the *vir* operon. Here, we show that, unlike other plasmid silencers that target promoters, Sfx cooperates with Rho factor to arrest transcription elongation. ChIP-seq reveals that despite sharing similar DNA motifs and a preference for negative supercoiling, Sfx and H-NS occupy distinct niches: Sfx binds weakly to the chromosome but is enriched on the R6K *vir* operon, from which H-NS is excluded. We hypothesize that this selective targeting is mediated by Sfx-*vir* interactions and phase separation. We show that Sfx binding to *vir* DNA critically depends on DNA topology but not on the target location. Our results suggest that Sfx phase separates with R6K to ensure its preferential recruitment to the plasmid DNA and forms stable nucleoprotein filaments that are impermeable to competitors. These findings reveal how histone-like proteins can partition the genome into distinct regulatory niches, a strategy likely mirrored across all life.

**GRAPHICAL ABSTRACT:** 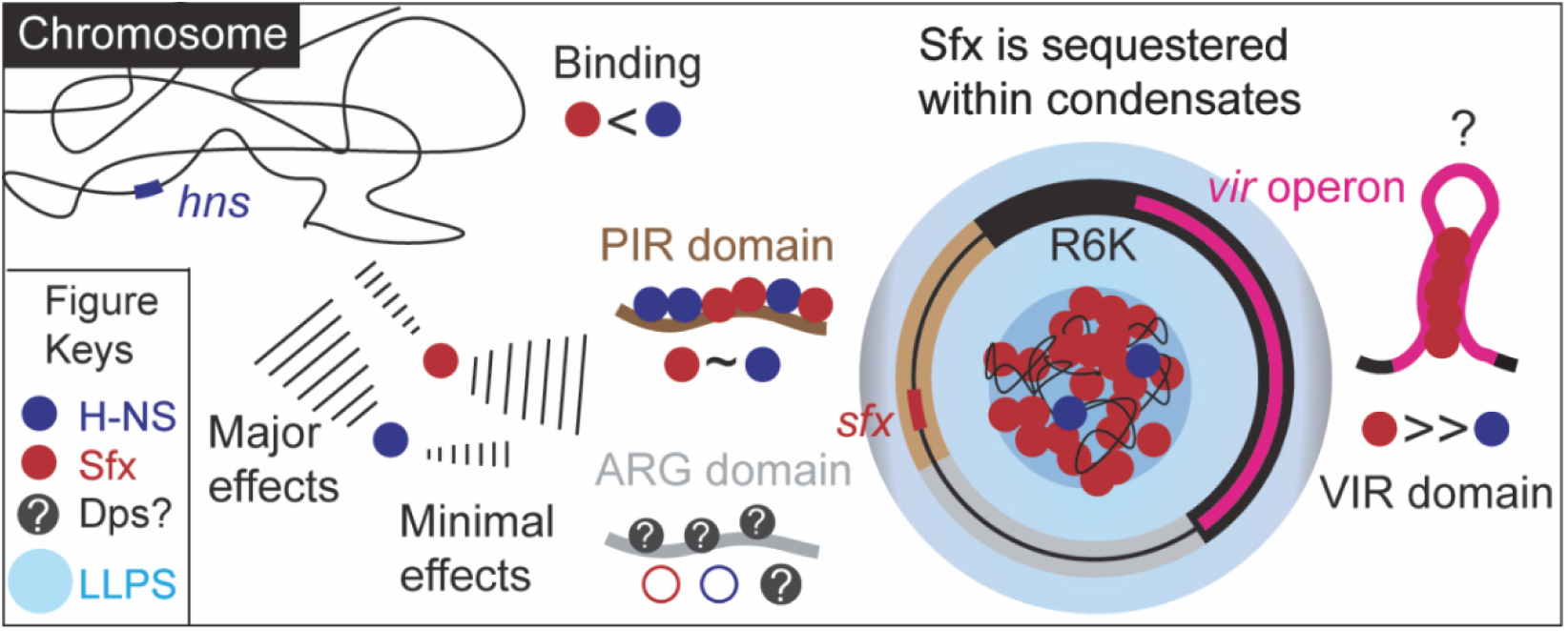

## INTRODUCTION

Conjugative plasmids autonomously transfer among bacteria, even across different species and genera (1), driving the bacterial evolution and the dissemination of multidrug resistance (2). Plasmids readily accumulate “beneficial” genes, such as those encoding resistance to antibiotics, toxins, and heavy metals, as well as virulence factors, and share these genes with other mobile genetic elements, facilitating the assembly of new combinations of traits (3).

While horizontal gene transfer enables rapid adaptation to hostile environments, it may come at a cost. Conjugative DNA transfer is mediated by large membrane-spanning type IV secretion system (T4SS) complexes (4) encoded by very long plasmid operons. Expression of the transfer operons (commonly called *tra* or *vir*; see (5) for the current classification) is associated with cellular stress (6) and can impose a metabolic burden (7). Even more crucially, the conjugative pili serve as receptors for phages (8). To minimize these risks, the *tra* genes must be tightly regulated, a process best understood in F (Supplementary Fig. S1), the first plasmid discovered in *Escherichia coli*; reviewed in (9).

Many long operons exhibit polarity, a decrease in the levels of RNAs transcribed from the distal genes of the operon. Polarity is imposed by Rho, a transcription termination factor that triggers premature release of RNAs that are not actively translated (10), a common feature of AT-rich xenogenes (11). Rho-dependent termination in xenogenes is aided by NusG, a ubiquitous transcription elongation factor that enables Rho to act in the absence of high-affinity, C-rich RNA elements (12,13), and H-NS, which binds to AT-rich DNA motifs and spreads to form extended nucleoprotein filaments (14). The H-NS filaments can hinder RNAP binding to promoters or create blocks to elongation (15), which are a prelude to Rho-dependent termination. Together, Rho, NusG, and H-NS silence pathogenicity islands and other xenogenes (16).

RfaH, a sequence-specific paralog of NusG, is recruited to the transcribing RNAP at the operon polarity suppressor (*ops*) DNA element (17,18) and relieves Rho and H-NS silencing. RfaH abolishes Rho-mediated polarity (19) by inhibiting RNAP pausing (18), excluding NusG from RNAP (20), and activating translation (21). RfaH also counter-silences H-NS (22,23), presumably by destabilizing bridged nucleoprotein filaments that are formed by H-NS and accessory proteins (StpA and Hha) to trigger premature termination (15).

As plasmids are mobile xenogenes, finding that they are regulated similarly to their sedentary relatives appears hardly surprising. However, some conjugative plasmids, including the well-studied R6K, encode paralogs of H-NS and RfaH, Sfx and ActX, respectively (Fig. 1A). R6K, the prototype plasmid of the IncX group, is a 40-kb IncX2 plasmid that has been used as a model system for plasmid replication and conjugation studies for decades. R6K contains three antibiotic-resistant genes located in a GC-rich transposition island, genes required for plasmid replication, segregation, and transfer (the *vir* operon and relaxosome *taxA/C* genes), and encodes three recently identified regulatory proteins, ActX, VirBR, and Sfx (Fig. 1A). VirBR and ActX have been reported to activate conjugation in other IncX plasmids (24,25), while Sfx silences R6K conjugation (26).

**Figure 1.**
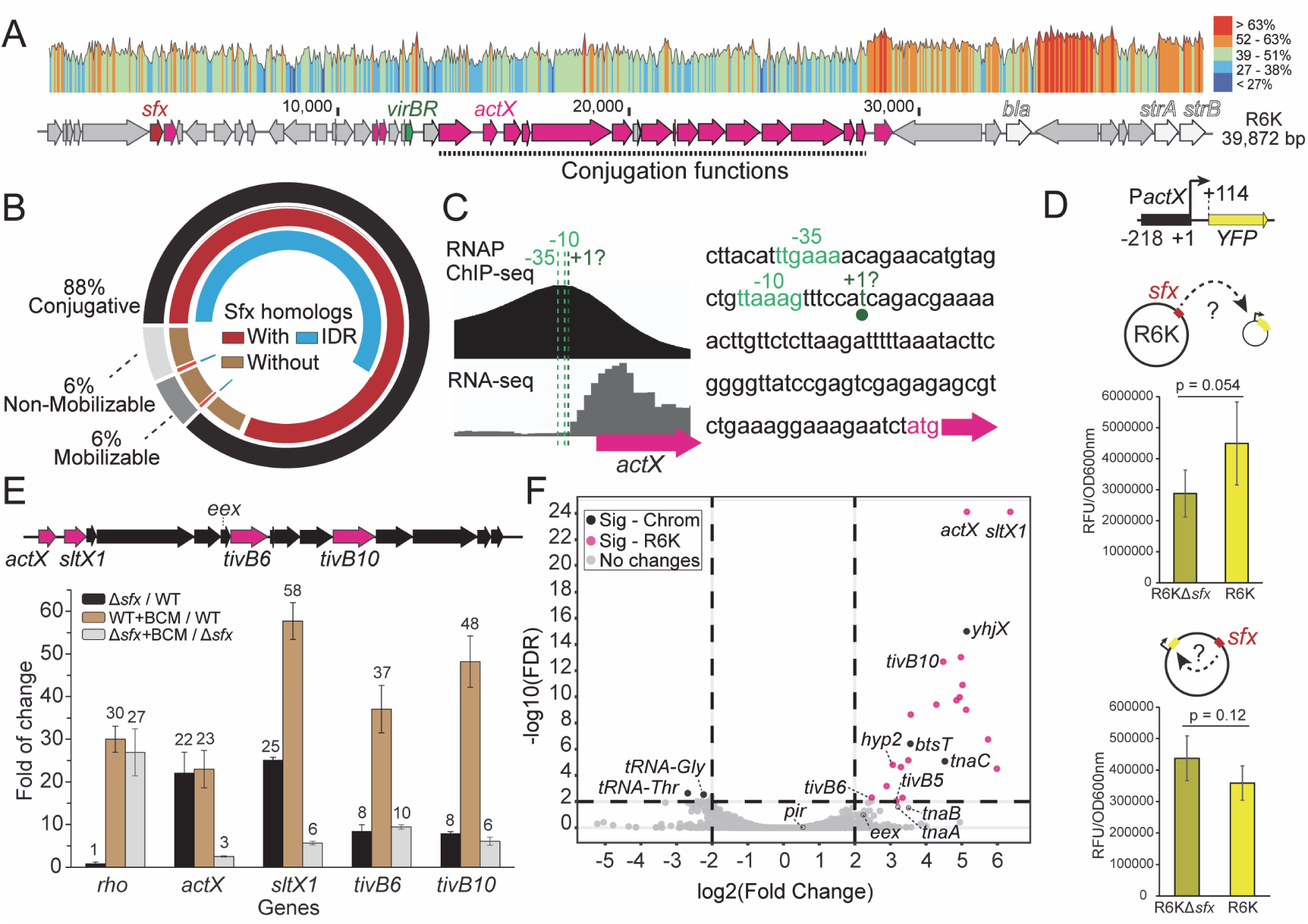
R6K encoded Sfx represses the plasmid conjugation functions. (**A**) R6K map. The three antibiotic-resistant genes (*bla*, *strA*, *strB*) are in light grey; genes that encode predicted positive regulators of conjugation, *virBR* (green) and *actX* (magenta), and the silencer *Sfx* (red) are indicated. The GC content (calculated with Snapgene v8.2.2) is shown above the map. Genes significantly upregulated by the *sfx* deletion are highlighted in magenta (see panel **F**). (**B**) Sfx is the core gene of IncX plasmids (a total of 1881 IncX plasmids were analyzed). The outer ring shows the distribution of the three mobility groups in IncX plasmids. The middle ring indicates the presence of Sfx homologs. The inner ring shows the presence of IDRs in Sfx homologs. (**C**) RNAP ChIP-seq reveals the location of P*actX* promoter. The -10 and -35 elements are predicted by BPROM, and the transcription start site (+1) is predicted by BDGP. RNA-seq track is smoothed in 20 bp windows. The sequences of -10 and -35 elements are shown on the right. (**D**) Relative fluorescence unit (RFU) of yellow fluorescent protein (YFP), normalized by OD_600_ nm. The effects of Sfx on P*actX* were tested using the P*actX*-*yfp* reporter located on a P15A plasmid (*in trans*, top) or on R6K (*in cis*, bottom). Error bars are the standard deviation (SD; n = 5). A two-tailed t-test assuming unequal variances was used to calculate p-values. (**E**) RT-qPCR analysis of selected genes. WT, wild-type R6K. BCM, bicyclomycin. Δ*sfx*, R6KΔ*sfx*. Mean values are indicated at the top of the bars. Error bars are SD (n = 3). (**F**) Volcano plot of RNA-seq data. The significantly changed genes (FDR < 0.01, ABS(log2FC) > 2, FC (Fold Change) = R6KΔ*sfx* / R6K WT) are highlighted in black for chromosomal and magenta for R6K genes.

Our bioinformatic analysis (Dataset 1) shows that 88% of IncX plasmids are predicted to be conjugative (Fig. 1B), and more than 90% of these encode one or two Sfx homologs. In comparison, only 7 – 8% of mobilizable or non-mobilizable plasmids have Sfx. Thus, *sfx* is a core gene in conjugative IncX plasmids that silences conjugation of R6K (26) and other IncX plasmids (24,25). A recent phylogenetic analysis reveals that Sfx has diverged from the H-NS paralog StpA, undergoing positive selection in several regions, including the key DNA-binding AT-hook motif (26). Similar to StpA (27) and Sfh, an H-NS like protein encoded on the IncHI1 plasmid (28,29), Sfx could partially complement the phenotypes of the *hns* deletion (26); however, H-NS did not inhibit conjugation of R6KΔ*sfx* (26). This one-way complementation is particularly striking given that H-NS is known to bind AT-rich DNA, such as the R6K *vir* operon (Fig. 1A), with loose specificity (reviewed in (11)). The inability of H-NS to substitute for Sfx suggests that H-NS may be excluded from the R6K *vir* operon to enable autonomous conjugation control. Alternatively, the binding of H-NS to the *vir* operon may not be able to silence transcription. However, nothing is known about the molecular details of Sfx binding to chromosomal and plasmid DNA, and the mechanism by which Sfx silences R6K conjugation.

Here, we show that Sfx shares many chromosomal targets with H-NS but exhibits an apparently reduced occupancy. Consistently, the *sfx* deletion does not alter transcription of most associated chromosomal genes. This pattern is reversed on R6K: Sfx binds to and strongly represses the *vir* operon during transcription elongation, acting together with Rho, whereas H-NS binds poorly to most R6K regions (except the *pir* gene, which encodes the R6K replication initiation protein) and has no effect on R6K conjugation. Our analysis of DNA recognition motifs, target shuffling, and *in vitro* DNA binding assays show that the selective recruitment of Sfx to DNA is mediated by sequence-specific contacts and is influenced by template topology but is less sensitive to the target location. Finally, we show that Sfx can form condensates with R6K, a property that may favor its preferential binding to the plasmid DNA.

## MATERIALS AND METHODS

Plasmids, oligonucleotides, and strains used in this study are listed in Supplementary Table S1. Restriction and modification enzymes for plasmid construction were obtained from New England Biolabs, DNA - from Millipore Sigma, and synthetic DNA fragments for HiFi assembly - from Genscript. The sequence of all plasmids was confirmed by whole plasmid sequencing at Plasmidsaurus; the GenBank format files are included in Dataset 2. Antibiotics were added when needed: carbenicillin (100 μg/ml), kanamycin (40 μg/ml), chloramphenicol (30 μg/ml), and tetracycline (15 μg/ml).

### Sfx homologs distribution in IncX plasmids

IncX plasmid metadata, including NCBI accession ID and predicted mobility, were downloaded from PLSDB (v. 2024_05_31_v2) (30). Multi-replicon plasmids were excluded from further analysis. A total of 1,881 IncX plasmid sequences were downloaded from NCBI. Using the R6K Sfx as a query, a BLASTp search (BLAST + v.2.9.0) (31) was performed with an E-value cutoff of 10^−10^ against the downloaded sequences to collect Sfx homologs. In all BLAST searches, truncated homologs that consist solely of an N- or C-terminal domain were excluded from consideration by restricting searches to proteins with > 50% coverage of Sfx. The intrinsically disordered regions (IDRs) were predicted by metapredict v3 (32).

### Protein purification

*E. coli* BL21 (DE3) cells were used for protein overexpression. Cells harboring the expression vector (Supplementary Table S1) were cultured in Terrific Broth (Research Products International, cat# T15100) at 37 °C, and isopropyl-1-thio-β-D-galactopyranoside (IPTG; Goldbio, cat# I2481C) was added when OD_600_ reached ∼ 0.5. For H-NS and Sfx, 0.5 mM IPTG was used to induce expression for 2.5 h at 37 °C. For Hha, 0.1 mM IPTG was used for an overnight induction at 16 ℃. The cells were collected by centrifugation at 8,000 × *g* for 8 min at 4 °C.

**H-NS**: the cell pellet was resuspended in Lysis buffer A (50 mM Tris-HCl, pH 8, 5% (v/v) glycerol, 1 M KCl, 10 mM β-mercaptoethanol (β-ME), 1 x ProBlock Gold 2D Protease Inhibitor Cocktail (EDTA Free; Goldbio, cat# GB-109-1)) and opened by sonication. The cell lysate was cleared by centrifugation (20,000 × *g*) for 30 min at 4 °C. The supernatant was passed through a 0.45 μm filter and applied to High Affinity Ni-charged Resin (Genscript, cat# L00223). The resin was washed with Ni-A buffer (40 mM Tris-HCl, pH 8, 5% glycerol, 1 M KCl, 10 mM β-ME, 0.2 mM phenylmethylsulfonyl fluoride (PMSF; ThermoFisher, cat# 36978), 40 mM imidazole), and protein was eluted with Ni-B buffer (40 mM Tris-HCl, pH 8, 5% glycerol, 50 mM KCl, 10 mM β-ME, 500 mM imidazole). The eluted protein was loaded onto the HiTrap Heparin HP column (Cytiva, cat# 17040701) and eluted with a linear gradient of KCl (0 – 1 M) in Hep buffer (20 mM Tris-HCl, pH 8, 5% glycerol, 1 mM tris(2-carboxyethyl)phosphine (TCEP)).

**Hha**: the cell pellet was resuspended in Lysis buffer B (100 mM HEPES-KOH, pH 7.5, 300 mM KCl, 5% glycerol, 1x cOmplete protease inhibitor cocktail (EDTA-free; Roche Diagnostics, cat# 11836170001], 10 mM imidazole, 5 mM β-ME), and lysed by sonication. The cleared lysate was applied to Ni^2+^-NTA resin (Cytiva, cat# 17531801), washed with Ni-C buffer (20 mM HEPES-KOH, pH 7.5, 300 mM KCl, 5% glycerol, 5 mM β-ME, 0.1 mM PMSF) supplemented with 30 mM imidazole, and eluted with Ni-buffer D (20 mM HEPES-KOH, pH 7.5, 50 mM KCl, 5% glycerol, 5 mM β-ME, 0.1 mM PMSF, 300 mM imidazole). The eluted protein was buffer exchanged to SEC buffer (20 mM HEPES-KOH, pH 7.5, 300 mM KCl, 5% glycerol, 5 mM β-ME) with a HiPrep 26/10 desalting column (Cytiva, cat# 17508701).

**Sfx**: the cell pellet was resuspended in Lysis buffer A and opened by sonication. The cell lysate was cleared by centrifugation (20,000 × g) for 30 min at 4 °C. The supernatant was passed through a 0.45 μm filter and applied to High Affinity Ni-charged Resin (Genscript). The resin was washed with Ni-A buffer, and protein was eluted with Ni-C buffer (40 mM Tris-HCl, pH 8, 5% glycerol, 0.5 mM KCl, 10 mM β-ME, 500 mM imidazole). The eluted protein was loaded onto a desalting column (Cytiva) in desalting buffer (40 mM Tris-HCl, pH 8, 5% glycerol, 1 mM TCEP, 0.4 M KCl) to remove imidazole.

His-SUMO-tagged H-NS, Hha, and Sfx proteins were treated with ULP1 protease (purified in-house) at 4 °C overnight to remove the tag. The sample was passed through Ni-charged resin again to remove the ULP1 protease and the tagged proteins. The untagged protein was loaded onto a Superdex 75 10/300 GL column (Cytiva, cat# 17517401). Protein purity was assessed by SDS-PAGE and Coomassie staining. Protein concentrations were determined by measuring the absorbance at 280 nm on a Nanodrop ND-1000 spectrometer. Purified proteins were aliquoted and stored at −80 °C.

### Protein labeling

A single cysteine residue was added to the C-terminus of H-NS and Sfx (Supplementary Table S1). The proteins were first purified as above without cleaving the His-SUMO tag, followed by on-bead labeling. The His-tagged proteins were first reduced by incubating with Ni-charged MagBeads (GenScript, cat# L00295) in desalting buffer for 3 hours at 4 °C on a rotating wheel. Then the magnetic beads were washed with PBS buffer (137 mM NaCl, 2.7 mM KCl, 8 mM Na_2_HPO_4_, and 2 mM KH_2_PO_4_, pH 7.4; degassed for half an hour before use) and resuspended in PBS. Fluorescent dyes were diluted 75 times into the resuspended beads. Alexa Fluor 546 C5 Maleimide (AF546; ThermoFisher, cat# A10258) and Cyanine 5 maleimide (Cy5; AAT Bioquest, cat# 152) were used for Sfx and H-NS derivatives, respectively. After overnight incubation at 4 °C, the beads were washed with desalting buffer and then treated with ULP1 for 4 hours at 4 °C to release the labeled proteins from magnetic beads. Protein purity was assessed by SDS-PAGE, and the protein concentration was determined by the Bradford protein assay. The proteins were aliquoted, flash frozen in liquid nitrogen, and stored at −80 °C.

### Reverse transcription quantitative PCR (RT-qPCR)

*E. coli* BW25113 carrying R6K or R6KΔ*sfx* were cultured in LB with carbenicillin, and *E. coli* MG1655 derivatives carrying F-factor (IA1016, IA1017, IA1018; Supplementary Table S1) were cultured in LB with chloramphenicol. The cells were grown at 37 °C to an OD_600_ of ∼0.7 and treated with bicyclomycin (BCM; 50 μg/mL) for 15 min, where indicated. The cells were mixed with 0.2 volume of ice-cold growth stop solution (5% phenol and 95% ethanol) to inactivate cellular RNase before being collected by centrifugation at 4,000 × g for 10 min. The mRNA was isolated using the Monarch Total RNA Miniprep Kit (NEB, cat# T2010S). RNA samples were treated with TURBO DNase (ThermoFisher, cat# AM2239) before RT-qPCR. A total of 100 ng RNA samples were used with iTaq Universal SYBR green one-step kit (Bio-Rad, cat#1725150) and analyzed on a CFX96 system (Bio-Rad). Samples without reverse transcriptase were used as a negative control to ensure the absence of DNA contamination. The quantification cycle (Cq) values were calculated using CFX Manager v3.0 in regression mode. The gene expression level was analyzed by the threshold cycle (2^−ΔΔCT^) method (33); the *ihfB* gene was used as a reference, expression of which is not affected by BCM (34).

### Next-generation sequencing (NGS) sample preparation

#### RNA-seq samples

*E. coli* BW25113 carrying R6K or R6KΔ*sfx* were grown in LB medium supplemented with carbenicillin at 37 °C and 250 rpm. Cells were collected by centrifugation when the OD_600_ reached 0.6. Total RNA was first extracted with RNAzol RT (Millipore Sigma, cat# R4533), and the top water phase was fed into Monarch Total RNA Miniprep Kit (NEB). The extracted RNA was further treated with DNase I (NEB, cat# M0303S). The rRNA was removed by using NEBNext rRNA Depletion Kit (Bacteria) (NEB, cat# E7850S) following the manufacturer’s protocol. The RNA samples were submitted for stranded RNA sequencing.

### Sfx and H-NS ChIP-seq samples

Chromatin immunoprecipitation followed by sequencing (ChIP-seq) was used to identify Sfx and H-NS binding sites. The procedure was adapted from a previous publication (35). FLAG tag was added to the C-terminus of H-NS and Sfx. The overnight cultures (∼16 hours) of cells containing either H-NS-3xFLAG encoded on the chromosome or Sfx-FLAG expressed from a low copy plasmid under its native promoter (p451) (Supplementary Table S1) were mixed with formaldehyde/sodium phosphate (pH 7.4) buffer to yield a final concentration of 10 mM NaPO_4_ and 1% (v/v) formaldehyde. The cells were crosslinked for 5 min at 37 °C, 250 rpm. The crosslinking was stopped by adding 0.33 M glycine, followed by a 30-min incubation in an ice-water bath. The crosslinked cells were collected by centrifugation, washed three times with PBS, and resuspended in 1x IP buffer (200 mM Tris-HCl, pH 8, 600 mM NaCl, 4% Triton X-100, 2x ProBlock Gold 2D Protease Inhibitor Cocktail (Goldbio)). The resuspended cells were subjected to controlled sonication to yield DNA fragments ranging from 100 to 500 bp. The cells were kept in an ice-water bath throughout sonication.

After sonication, 0.1 volume of cell lysate, which is used as the input sample, was kept at 4 °C until the reversal of crosslinking. The remaining cell lysate (ChIP sample) was mixed with MonoRab anti-DYKDDDDK magnetic beads (GenScript, cat# L00835-1). After an overnight incubation at 4 °C, the magnetic beads were subjected to sequential washes, with 1 ml used per wash: 1) 1x Wash buffer A (100 mM Tris-HCl, pH 8, 250 mM LiCl, 2% Triton X-100, 1 mM EDTA); 2) 1x Wash buffer B (100 mM Tris-HCl, pH 8, 500 mM NaCl, 1% Triton X-100, 0.1% sodium deoxycholate, 1 mM EDTA); 3) 1x Wash buffer C (10 mM Tris-HCl, pH 8, 500 mM NaCl, 1% Triton X-100, 1 mM EDTA); 4) 1x TE (10 mM Tris-HCl, pH 8, 1 mM EDTA). The antigens were eluted in 1 ml of elution buffer (50 mM Tris-HCl, pH 8; 10 mM EDTA; 1% SDS, 5 U/ml proteinase K (NEB, cat# P8107S)) during 1 h-incubation at 65 °C, with vigorous vortexing every 10 minutes. The input samples were diluted 10 times with elution buffer and subjected to the same treatment as the ChIP samples from now on. After removing the magnetic beads from ChIP samples, both ChIP and input samples were incubated at 65 °C overnight to reverse the crosslinking. To digest the RNA, 100 μg/ml of RNase A (ThermoFisher, cat# EN0531) was added for a 2 h incubation at 37 °C, followed by 2 h incubation at 50 °C with 5 U/ml of proteinase K (NEB). Finally, DNA was recovered using QIAquick PCR Purification Kit (QIAGEN, cat# 28104). Recovered DNA samples were submitted for sequencing.

### RNAP ChIP-seq samples

*E. coli* MG1655 cells harboring R6KΔ*sfx* were grown in LB at 37 °C, 250 rpm, and were treated with 150 μg/ml rifampicin for 10 min at OD_600_ ∼0.4. The treated cells were crosslinked and washed as above. The protocol of immunoprecipitation is largely the same as Sfx ChIP-seq sample preparation except for the incubation with antibodies. The samples were first incubated with Protein A/G MagBeads (GenScript, cat# L00277) for 3 hours at 4 °C, and the protein A/G MagBeads were then removed. Anti-*E. coli* RNA polymerase β subunit antibody (abcam, cat# ab191598) was added to the samples for an overnight incubation at 4 °C. The next day, Protein A/G MagBeads were added again. After 3 hours of incubation, the magnetic beads were washed, and the DNA was extracted as above.

### RNA ChIP-seq samples

The protocol is largely the same as Sfx ChIP-seq sample preparation, with the following changes. After reverse crosslinking, the samples were treated with Turbo DNase (ThermoFisher) instead of RNase A, and the RNA was purified by Monarch Total RNA Miniprep Kit (NEB) following the manufacturer’s protocol. The resulting RNA was submitted for stranded RNA sequencing.

All samples were submitted to IDI-GEMS laboratory, The Ohio State University, for NGS. To generate libraries for DNA samples, the Illumina DNA Prep kit (Illumina, cat# 20060060) was used following the procedure for selecting < 500 bp fragments. Samples were barcoded using IDT for Illumina UD Indexes. Stranded Total RNA Library Prep (Illumina, cat#20040525) was used for RNA samples following Illumina’s protocol. Library concentration was measured using a Qubit Flex fluorometer (ThermoFisher), and the library fragment size was analyzed using a TapeStation 4150 (Agilent).

### NGS data analysis

#### RNA-seq

Adapter and low-quality reads were trimmed using TrimGalore v0.6.7 (36) in paired-end mode and one base was trimmed from the 5’ end of all reads (Parameters: --paired --trim-n-clip_R1 1 --clip_R2 1). Reads were aligned using bowtie2 v2.4.4 (37) with very sensitive, end-to-end alignment, and dovetailed alignments allowed (Parameters: --very-sensitive --no-mixed --no-discordant --dovetail -X 1000). The reads were mapped to *E. coli* K-12 substr. BW25113 (ASM75055v1) and plasmid R6KΔ*sfx* (pIA1548; Dataset 2). The R6K sequence used in this study is the same as NCBI ID LT827129.1, except that we shifted the coordinates to group the high GC-region for visualization (Fig. 1A, Dataset 2). The alignment files were converted to BAM format and indexed by SAMtools v1.23 (38). The alignment files were subjected to quantification using featureCounts v2.0.3 (39) with the --countReadPairs option. Finally, the differential expression was analyzed using DEseq2 v3.22 (40). The IGV tracks were generated using deepTools v3.5.6 (41).

#### ChIP-seq

Adapter and low-quality reads were trimmed using TrimGalore v0.6.7 (36) in paired-end mode (Parameters: --paired --trim-n), and “--2colour 30” was added to the parameters if poly-G artifacts were present in the reads. For RNA ChIP-seq reads, parameters “--trim-n -q 28 --clip_R1 1 --clip_R2 1 --paired” were used. Reads were aligned using bowtie2 v2.4.4 (37) with very sensitive, end-to-end alignment, and dovetailed alignments allowed (Parameters: --very-sensitive --no-mixed --no-discordant --dovetail -X 1000). The alignment files were converted to BAM format and indexed by SAMtools v1.23 (38). Peak calling was done by MACS3 v3.0.3 (42) with q cutoff < 10^-7^ to remove peaks with fold enrichment (FE = ChIP/Input) < 1.5. Searching for binding motifs of H-NS and Sfx was done by HOMER v5.1 (43) using peaks on the chromosome. The FE tracks displayed in IGV v2.19.7 (44) were calculated by MACS3 and converted to bigWig files by bedGraphToBigWig v2.10 (45). The -10 and -35 elements of promoters were predicted by BPROM (46), and the transcription start site was predicted by BDGP (47). Searching the high-affinity binding sites (named Seeds) of Sfx on R6K was done by Fuzzy Search DNA (48) with the identified motifs in this study.

### ChIP-qPCR

The samples were prepared as Sfx and H-NS ChIP-seq sample preparation. The resulting DNA was submitted for qPCR instead of NGS. The qPCR was performed on a CFX96 system (Bio-Rad) with iTaq Universal SYBR Green Supermix (Bio-Rad, cat# 1725120). Primers for qPCR are listed in Supplementary Table S1. One ng of DNA was loaded into each well. The Cq values were calculated by CFX Manager v3.0 using regression mode. The fold enrichment of the target fragment was analyzed by the threshold cycle (2^−ΔΔCT^) method (33).

### Electrophoretic mobility shift assay (EMSA)

Linear DNA templates were generated by PCR amplification (Supplementary Table S1) and purified using QIAquick PCR purification kit (QIAGEN). SsrA RNA was generated by *in vitro* transcription with T7 RNA polymerase P266L mutant (49). The reaction in 40 mM Tris-HCl, pH 7.9, 10 mM NaCl, 6 mM MgCl_2_, 10 mM DTT, 2 mM spermidine, 0.05% Tween 20, and 0.5 mM of each nucleoside triphosphate (ATP, CTP, GTP, UTP) was incubated at 30 °C for 2.5 h. After treatment with TURBO DNase (Thermofisher), RNA was purified by Monarch Spin RNA Cleanup Kit (NEB, cat# T2030S). Radioactive labeling of DNA and RNA templates was done using T4 PNK (NEB) and ATP [γ-^32^P] (Revvity, cat# BLU002Z).

Indicated protein and template were incubated in EMSA buffer (25 mM Tris-HCl, pH 7.5, 100 mM KCl, 5 mM MgCl_2_, and 1 mM TCEP) at room temperature (∼ 23 °C) for 15 min. Then the reactions were mixed with 20× loading buffer (30% glycerol, 0.2% Orange G) and separated on 3% polyacrylamide or 1% agarose gels in 0.5× TBE (Research Products International, cat# T22020) at 4 °C. The DNA and RNA species were visualized by Amersham Typhoon 5 and quantified by Fiji v2.17.0 (50).

### Pelleting assay

Reactions containing different protein, plasmid, and RNA combinations were assembled in 20 μl EMSA buffer. After 15 min incubation at room temperature, the reactions were centrifuged at 21,000 × *g* for 25 min at 20 °C. The supernatant was removed, and the pellet was soaked in 15 μl of HSB buffer (20 mM Tris-HCl, pH7.5, 5 mM DTT, 500 mM NaCl) and 5 μl of 4x LDS sample buffer (GenScript, cat# M00676) for 30 min at room temperature before resuspending. The samples were analyzed on SurePAGE 4-12% gels (GenScript, cat# M00654) and stained with GelCode Blue Safe Protein Stain (ThermoFisher, cat# 24594). The gels were visualized by Amersham Typhoon 5 and quantified by Fiji v2.17.0 (50).

### Plasmid copy number determination

Plasmid copy number was determined by a qPCR-based method modified from a previously published protocol (51). The *rpoC*, *bla*, and *cat* genes were selected as target genes for the chromosome, R6K, and p451, respectively. The amplification efficiencies of primers of *rpoC*, *bla*, and *cat* are determined to be ∼1 (Supplementary Table S1). About 10^9^ cells were collected by centrifugation and resuspended in 100 μl of 20 mM Tris-HCl, pH 8, 20 mM EDTA, 0.2% SDS. After incubating for 5 min at 95 °C to open the cells, RNase A (ThermoFisher) was added to a final concentration of 0.2 μg/μl. The reaction was incubated for 1 h at 37 °C, followed by incubation with 5 U/ml Protease K (NEB) for 1 hour at 50 °C. Finally, the sample was incubated for 10 min at 95 °C and centrifuged at 4 °C, 21,000 × *g* for 10 min to pellet insoluble fibers. The cleared supernatant was diluted 500 times with DNase-free H_2_O, and 1 μl of the diluted sample was used for qPCR. The iTaq Universal SYBR Green Supermix (Bio-Rad) was used for qPCR analysis on the CFX96 system. The plasmid copy number is expressed as the ratio of plasmid gene copy to chromosomal gene copy, which was calculated using the threshold cycle (2^−ΔΔCT^) method (33).

### Conjugation assay

*E. coli* BW25113 strains harboring R6K derivatives were used as donors, and *E. coli* BW25113 *lysA*::Kn^R^ strain was used as the recipient. Donors and recipients were grown overnight in LB supplemented with selective antibiotics at 37 °C, 250 rpm. Overnight cells were collected by centrifugation, and the pellet was washed once with sterile PBS. Following the wash, 10^9^ donor cells and 10^9^ recipient cells were mixed and dropped onto an MCE membrane filter (0.22 µm pore size; Millipore Sigma, cat# GSWP01300) overlaid on LB agar and incubated at 30 °C for 2 h. The filter paper was then vortexed in 1 ml of sterile PBS. Serial dilutions of each sample were spotted on M9 (ATCC Medium 2511) + carbenicillin, LB + kanamycin, and LB + kanamycin + carbenicillin to select for donor, recipient, and transconjugant populations, respectively.

For testing the effects of novobiocin on conjugation, *E. coli* BW25113 *argE*::tet^R^ carrying R6K75 (a transposon-deletion R6K; Supplementary Table S1, Dataset 2) or R6K75Δ*sfx* were used as donors, and *E. coli* BW25113 *lysA*::Kn^R^ was the recipient. The overnight cells were reinoculated into fresh medium. At an OD_600_ of 0.4, cultures were exposed to novobiocin (250 μg/ml) for one hour, followed by washing once with sterile PBS. 500 μl of culture mixtures in PBS containing 10^9^ donor cells and 10^9^ recipient cells were incubated at 37 °C for 30 min, then vortexed to stop conjugation. Serial dilutions of each sample were then spotted on LB + tetracycline + carbenicillin, LB + kanamycin, and LB + kanamycin + carbenicillin to select for donor, recipient, and transconjugant populations, respectively. Plates were incubated overnight at 37 °C.

### Western blot

*E. coli* MG1655 carrying R6K75*sfx*-FLAG (pIA1721; Supplementary Table S1) and *E. coli* MG1655 *hns*-3xFLAG carrying R6K75 were cultured overnight in LB + carbenicillin, and the cells were collected by centrifugation. The cells were opened by sonication in 25 mM Tris-HCl, pH 7.4, 7 M Urea, and followed by resolving in 4 – 12% SurePAGE gel (GenScript). Then, the proteins were transferred to a nitrocellulose membrane (Bio-Rad, cat# 1620112) in a Trans-Blot SD Semi-Dry Transfer Cell (Bio-Rad) with 15 V for 15 min in Transfer buffer (25 mM Tris-HCl, pH 8.3, 192 mM glycine, and 20% methanol). After the transfer step, the membrane was blocked in Blocking buffer (25 mM Tris-HCl, pH 7.4, 150 mM NaCl, 0.1% Tween-20, and 5% (w/v) Blotting Grade Blocker Non-Fat Dry Milk (Bio-Rad, cat# 706404XTU)) for 1 h at room temperature. The membrane was washed twice with TBST (25 mM Tris-HCl, pH 7.4, 150 mM NaCl, 0.1% Tween-20) followed by incubation with anti-FLAG polyclonal antibodies (1:10,000) (Millipore Sigma, cat# F7425) in Blocking buffer overnight at 4 °C. The next day, the membrane was washed five times with TBST and incubated with secondary antibody (1:10,000; Goat Anti-Rabbit IgG (H + L)-HRP Conjugate (Bio-Rad, cat# 1706515)) in Blocking buffer for 1 h at room temperature. The membrane was then washed five times with TBST. Finally, the membrane was incubated with Clarity Max Western ECL Substrate (Bio-Rad) for 5 min at room temperature and imaged with ChemiDoc XRS + System (Bio-Rad). Quantification was done with Image Lab v6.1 (Bio-Rad).

### Yellow fluorescence protein (YFP) activity assay

*E. coli* BW25113 was used for the YFP activity assay. To test the effects of Sfx on the *actX* promoter (P*actX*), a YFP reporter was constructed by putting the *yfp* gene directly downstream of P*actX* (pIA1723; Supplementary Table S1). In order to test the *in trans* effects of Sfx, the P*actX*-*yfp* reporter was transformed into *E. coli* together with R6K or R6KΔ*sfx*. To establish the background, a control plasmid pBAD33 (p28; Supplementary Table S1), conferring the same chloramphenicol resistance as pIA1723, was transformed into *E. coli* together with R6K or R6K*Δsfx*. Five biological replicates of each strain were grown overnight in LB supplemented with carbenicillin and chloramphenicol at 37 °C, 250 rpm. The next morning, 1 μl of each replicate was used to inoculate 150 μl fresh media on a sterile, black-bottom 96-well plate (Greiner Bio-One, cat# 655077). Cells were allowed to grow for 24 hours at 37 °C with double-orbital shaking at 250 rpm in the VANTAstar plate reader (BMG). YFP fluorescence signal reads were taken every 15 minutes, and the final read was used for analysis. OD_600_ was used to normalize the fluorescence results.

To test the effects of Sfx *in cis*, P*actX*-*yfp* reporter was inserted into R6K or R6KΔsfx (pIA1767/1768; Supplementary Table S1). *E. coli* carrying R6K and *E. coli* carrying R6KΔsfx were used as background control. Cells were grown in LB supplied with carbenicillin. The testing procedure is the same as above.

### Atomic force microscopy

Shortly before deposition of the sample, a 5 μl droplet of 0.01 ug/ml of poly-L-ornithine (1 kDa MW, Millipore Sigma) was deposited onto freshly cleaved mica and incubated for 2 min. The poly-L-ornithine-coated mica was rinsed drop-wise with 400 μl of Nanopure (18 Mohm) water and dried gently with a stream of air from a portable compressor. Then, 5 μl of the sample solution containing 0.4 nM DNA plasmids was deposited on the poly-L-ornithine-coated mica and incubated for 2 min. This droplet was rinsed with 400 μl of Nanopure water and dried gently with compressed air. Images were acquired with a NanoScope MultiMode VIII AFM microscope (Bruker Nano Surfaces) operating in Tapping Mode using MicroMasch NSC:18 cantilevers with 5 nm nominal tip radius. Areas of 2 X 2 μm were scanned at a rate of ∼0.25 Hz with a resolution of 1024 x 1024 pixels. Images were filtered to remove scan line offsets and tilt/bow.

### Microscopy

Microscopy−based experiments were carried out in a Metamorph-driven Olympus BX-61. Glass slides (1 mm thickness; ThermoFisher, cat# 125444) and coverslips (0.17 mm thickness; VWR, cat# 48366-249) were washed with milliQ water (18 Mohm) and 70% ethanol, and dried in a 42 °C oven overnight. The dried glass slides and coverslips were coated with 1% (w/v) Pluronic F-127 (PF127; Millipore Sigma, cat# P2443). PF127 was prepared freshly in milliQ water. The treated slides and coverslips were rinsed gently with milliQ water and allowed to air-dry for 30 min. The proteins and R6K/RNA were mixed in a 10 μl reaction in EMSA buffer supplemented with polyethylene glycol 8000 (PEG8000). After 10 min at room temperature, the whole 10 μl was dropped onto the PF127-coated glass slide and covered with a PF127-coated coverslip. The droplets were visualized using a 40X objective in either bright field or fluorescence modes using an appropriate fluorescence channel.

## RESULTS

### Sfx promotes early termination in the R6K vir operon but has little effect on chromosomal genes

Conjugation is a fundamentally important process that drives horizontal gene transfer. Although R6K has been used as a conjugation model for decades (52), the promoters of its *vir* operon are still unknown. To understand R6K conjugation regulation, we first used RNAP ChIP-seq to identify potential promoters in R6K (Supplementary Fig. S2) upon treatment with rifampicin, an RNAP inhibitor that traps RNAP at promoters (53). Potential promoters were identified, for example, upstream from *pir*, *virBR*, and antibiotic resistance genes. Previous studies of IncX plasmids focused on the regulation of the major promoter of the conjugal transfer operon, demonstrating that its activity is controlled during initiation (54,55). Our ChIP-seq analysis revealed an RNAP peak upstream of the *actX* gene, P*actX*, an observation also supported by our RNA-seq data (Fig. 1C) and *in silico* promoter prediction.

To determine if Sfx inhibits RNAP recruitment to P*actX*, we constructed reporters in which the P*actX* promoter (-218 to +1) was fused to the *yfp* gene and tested in the presence (or absence) of the R6K-encoded Sfx (Fig. 1D). We compared the effects of Sfx expressed from R6K either *in trans,* with the reporter on a P15A plasmid (Fig. 1D top), or *in cis* on R6K (Fig. 1D bottom). In both cases, we observed robust YFP activity, which was not suppressed by Sfx, arguing that Sfx controls the *vir* operon during elongation (Supplementary Fig. S3AB). In fact, this mode of action could be expected because long xenogeneic operons such as the *vir* operon (∼13 kb) are frequently silenced by the termination factor Rho, which can act together with H-NS to induce premature RNA release during transcription of the chromosomal targets (12,56–58).

To evaluate this possibility, we tested the effects of Sfx and Rho on *vir* operon expression using RT-qPCR. Deletion of the *sfx* gene led to the upregulation of the four selected *vir* genes, whereas the chromosomal *rho* gene was not affected (Fig. 1E). Inhibition of Rho activity by bicyclomycin (BCM) increased the *rho* RNA levels 30 times, consistent with the autoregulation of the *rho* gene (59), as well as upregulation of the *vir* operon (Fig. 1E). Deleting *sfx* decreased the effects of BCM treatment (Fig. 1E). We conclude that Rho and Sfx synergistically repress the R6K conjugation operon.

This finding prompted us to investigate whether the F plasmid *tra* operon, commonly thought to be regulated during initiation (9), is also controlled during elongation. We found that Rho and H-NS silence the *tra* operon expression, whereas the transcription antitermination factor RfaH opposes their effects (Supplementary Fig. S1), a balancing act observed on RfaH-controlled other horizontally-acquired operons (22,23). Our results, along with the presence of genes that encode RfaH homologs on many conjugative plasmids, including R6K, suggest that transcription elongation control is a common regulatory strategy among these plasmids.

Some plasmid-encoded H-NS homologs have been reported to act as back-up copies of H-NS; for example, Sfh, encoded by the *Shigella flexneri* IncHI1 plasmid, shares the chromosomal binding sites and effects on gene expression with H-NS (28,29). To identify R6K and host genes controlled by Sfx, we performed RNA sequencing. Our results show that only a handful of genes are significantly affected by the *sfx* deletion (Fig. 1F, Dataset 1). Among these genes, only two tRNA genes located in the same *E. coli* operon were modestly downregulated, whereas all other genes were upregulated, consistent with the silencing role of Sfx. On R6K, most of these upregulated genes encode conjugation functions (Fig. 1AF). On the chromosome, three genes (*yhjX*, *btsT*, and *tnaC)* were significantly upregulated (Fig. 1F), all of which are implicated in pyruvate metabolism. The *tnaAB* operon leader peptide *tnaC* is significantly upregulated, but changes of *tnaA and tnaB* are not significant enough (FDR = 0.03), although they stand out from the chromosomal genes (Fig. 1F). Studies of ColE1 plasmid maintenance suggested that indole produced by the *tnaCAB* operon can inhibit cell division and plasmid replication to allow for multimer resolution (60). We speculate that the upregulation of *tnaCAB* increases pyruvate and indole production (61,62), which in turn could activate the BtsS/BtsR system, inducing *btsT* and *yhjX* to manage pyruvate homeostasis (63–65). Thus, *btsT* and *yhjX* changes might be caused by the *tnaCAB* operon.

Our results demonstrate that Sfx selectively silences the plasmid genes, whereas the published data show that H-NS controls many chromosomal genes (66), but does not inhibit R6K conjugation (67). To understand how these homologous silencers are directed to their respective regulons, we next determined the binding sites of Sfx and H-NS.

### Sfx weakly binds to many H-NS loci on the chromosome

We used ChIP-seq to analyze the genome-wide binding profile of H-NS and Sfx (Fig. 2A). H-NS is tagged with 3xFLAG, which has been widely used, and the tag does not substantially perturb H-NS function (68,69), whereas Sfx carries a single FLAG tag. This choice is based on our findings that Sfx’s ability to silence conjugation is sensitive to the presence of tags. The Sfx tagged with 1xFLAG (Sfx-FLAG) is fully active in silencing R6K conjugation, as previously shown (67) and confirmed in our experimental set-up (Supplementary Fig. S3AB). The Sfx-FLAG is expressed from its native promoter on a pSC101-derived plasmid (referred to as p451; Supplementary Table S1). We have determined the copy numbers of plasmids in the stationary phase, conditions used for conjugation and ChIP-seq assay (Fig. 2B): p451 is present at 3 – 4 copies/cell, and R6K – at 16 copies/cell; the latter number is within the published range (70). Our finding that Sfx can fully silence conjugation when expressed from the ∼4-fold less abundant locus (Fig. 2B, Supplementary Fig. S3B) suggests that Sfx expression may be autoregulated, as reported for H-NS (71).

**Figure 2.**
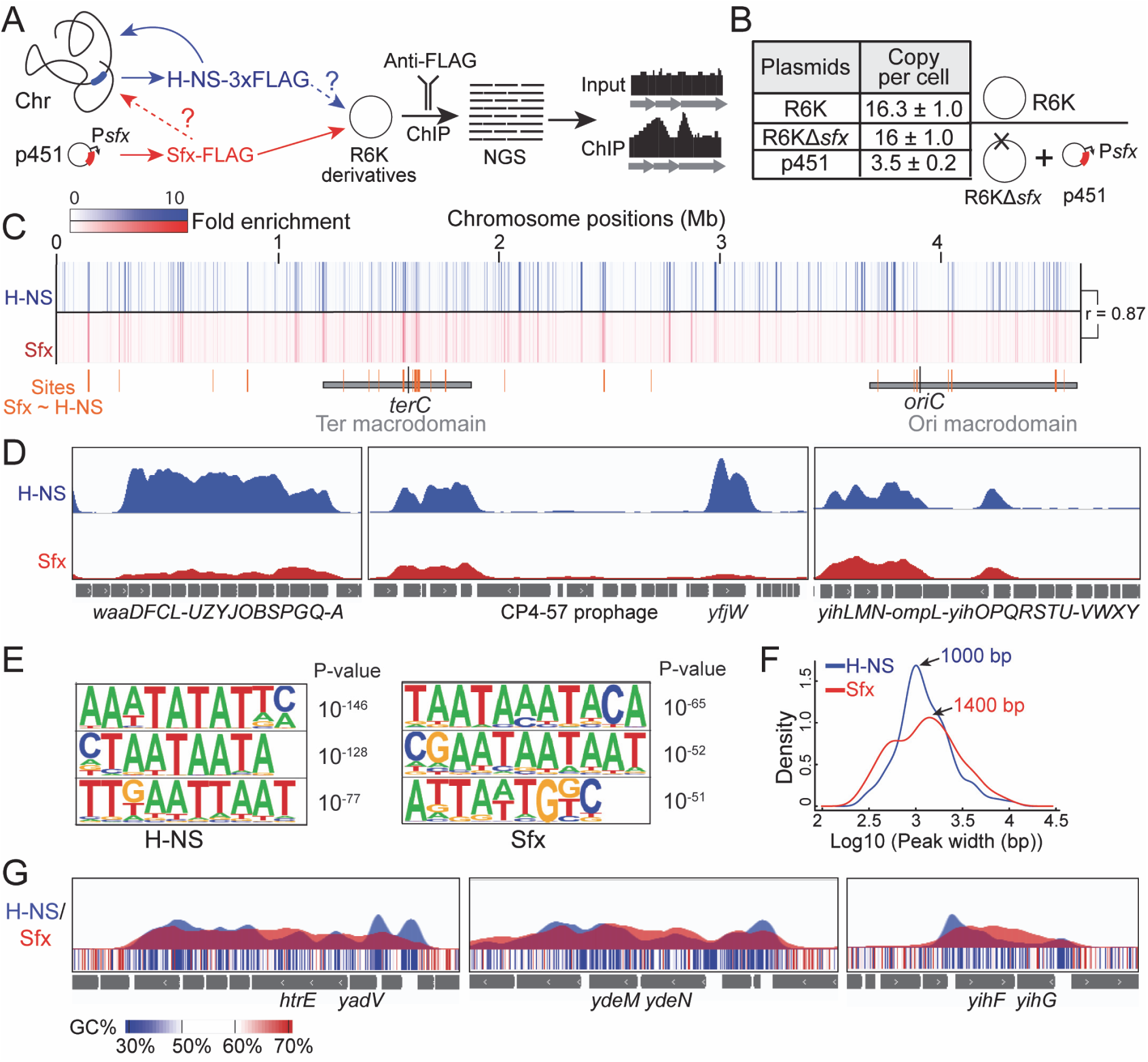
Sfx and H-NS share binding sites on the chromosome. (**A**) Schematic of the H-NS and Sfx ChIP-seq assay. Chr, chromosome. NGS, next-generation sequencing. (**B**) Plasmid copy numbers are shown as mean ± SD (n = 5). (**C**) Heatmap tracks of chromosome binding sites. Spearman correlation (r) analysis was performed on reads binned at 300 bp. Below the tracks, the two vertical black bars represent the replication terminus (*terC*) and origin (*oriC*). Orange bars are sites showing similar Sfx and H-NS binding, defined as fold enrichment (FE) at the peaks of Sfx >= 70% of H-NS peaks. Two horizontal grey bars are the chromosomal macrodomains. (**D**) Examples of H-NS and Sfx binding sites. The peaks in tracks represent fold enrichment (FE), which ranged from 0 to 13. (**E**) The binding motifs of Sfx and H-NS were predicted from the peaks on the chromosome by HOMER. The top 3 motifs with the lowest p-values are shown. (**F**) Density plot of the log10(width) of H-NS and Sfx peaks on the chromosome. The peak widths of the highest density are indicated. (**G**) Examples showing Sfx peaks extending through the valleys of H-NS peaks. The GC content, calculated in a 25 bp sliding window, is shown as a heatmap. The peaks in tracks represent FE, which ranged from 0 to 13.

On the chromosome, peaks of Sfx and H-NS largely overlap (with Spearman correlation of 0.87), though H-NS has higher overall occupancy at almost all loci (Fig. 2C). For example, H-NS has better occupancy than Sfx on the LPS core biosynthesis *waa* operon and *yfjW* gene of the CP4-57 prophage (Fig. 2D). The sites that show similar Sfx and H-NS peaks are mainly located in the replication terminus and origin macrodomains (Fig. 2C); see the *yih* operon for an example of similar binding (Fig. 2D). We note that these data cannot be used to compare true occupancies of the two proteins due to differences in their cellular concentrations, immunoprecipitation efficiency, and formaldehyde crosslinking between the proteins and across different DNA regions, a common limitation in ChIP studies. In addition, some of the observed shared sites may represent mixed filaments, which H-NS has been shown to form with its other homologs (15).

To estimate the relative levels of H-NS and Sfx under the conditions used for the ChIP-seq analysis, we used western blotting with anti-FLAG antibodies. Our results suggest that the molar ratio of Sfx/H-NS is at least 3.5 (Supplementary Fig. S3C), even though Sfx is expressed from a lower copy number plasmid (Fig. 2B). The higher abundance of Sfx is expected to be offset by the reduced affinity for the anti-FLAG antibodies; the manufacturer’s technical notes state that the 3xFLAG increases the detection limit in western blots by at least 10-fold compared to 1xFLAG. If this preference persists during immunoprecipitation, the observed H-NS/Sfx signal ratio might be overestimated by approximately 3-fold.

Selective protein targeting may be achieved through recognition of distinct DNA elements. Numerous studies of H-NS interactions with the DNA support the notion that H-NS prefers AT-rich sequences with loose specificity (72,73); following its loading at a high-affinity binding site, H-NS cooperatively spreads to adjacent regions (11,74,75). The majority of Sfx and H-NS peaks on the chromosome appear to coincide (Fig. 2C), suggesting that Sfx may recognize similar DNA motifs. We analyzed the Sfx and H-NS chromosomal peaks to compare their binding motifs; the top three motifs shown in Fig. 2E reveal that both proteins prefer AT-rich DNA, and their binding motifs can be part of a shared pattern AATAATAAATATAT (Fig. 2E). Interestingly, Sfx appears to tolerate more GC residues, and this property may partially explain its tendency to spread along the DNA in a more uniform pattern (see below).

Sfx peaks are generally broader than those of H-NS. While H-NS displays a maximum probability at 1000 bp peak width, Sfx peaks at 1400 bp and exhibits a higher probability of reaching widths beyond 3000 bp (Fig. 2F). Sfx also forms narrower peaks, which could be explained by the low fold enrichment. The peak boundary is determined by MACS3 (76) based on p-values, and a low fold enrichment can result in a non-significant p-value, pushing the boundary toward the peak center and producing narrower peaks. The wider peaks of Sfx are the result of shallow peak valleys. For example, chromosomal regions shown in Fig. 2G contain multiple H-NS peaks separated by deep valleys, whereas Sfx peaks are connected by shallow valleys and thus appear broadened. The same pattern is observed genome-wide.

### Sfx and H-NS have similar preferences for DNA topology

Since H-NS is known to preferentially bind to negatively supercoiled DNA (77–81), similar ChIP-seq peak locations (Fig. 2C) led us to assume that Sfx shares this preference. To further evaluate this premise, we compared H-NS and Sfx binding profiles to those of psoralen (82) and GapR (83) (Fig. 3A). Psoralen and GapR selectively bind to negatively- and positively-supercoiled DNA, respectively (83,84). Our analysis revealed that H-NS and Sfx profiles were similar to that of psoralen, yet quite distinct from the GapR profile (Fig. 3A), supporting the conclusion that both H-NS and Sfx prefer negatively supercoiled DNA. Consistently, the H-NS and Sfx profiles have a positive Pearson correlation with psoralen, but a negative correlation with GapR (Fig. 3B). We note that both positive and negative correlations were more pronounced with Sfx, suggesting that its preference for the negatively supercoiled DNA may be stronger.

**Figure 3.**
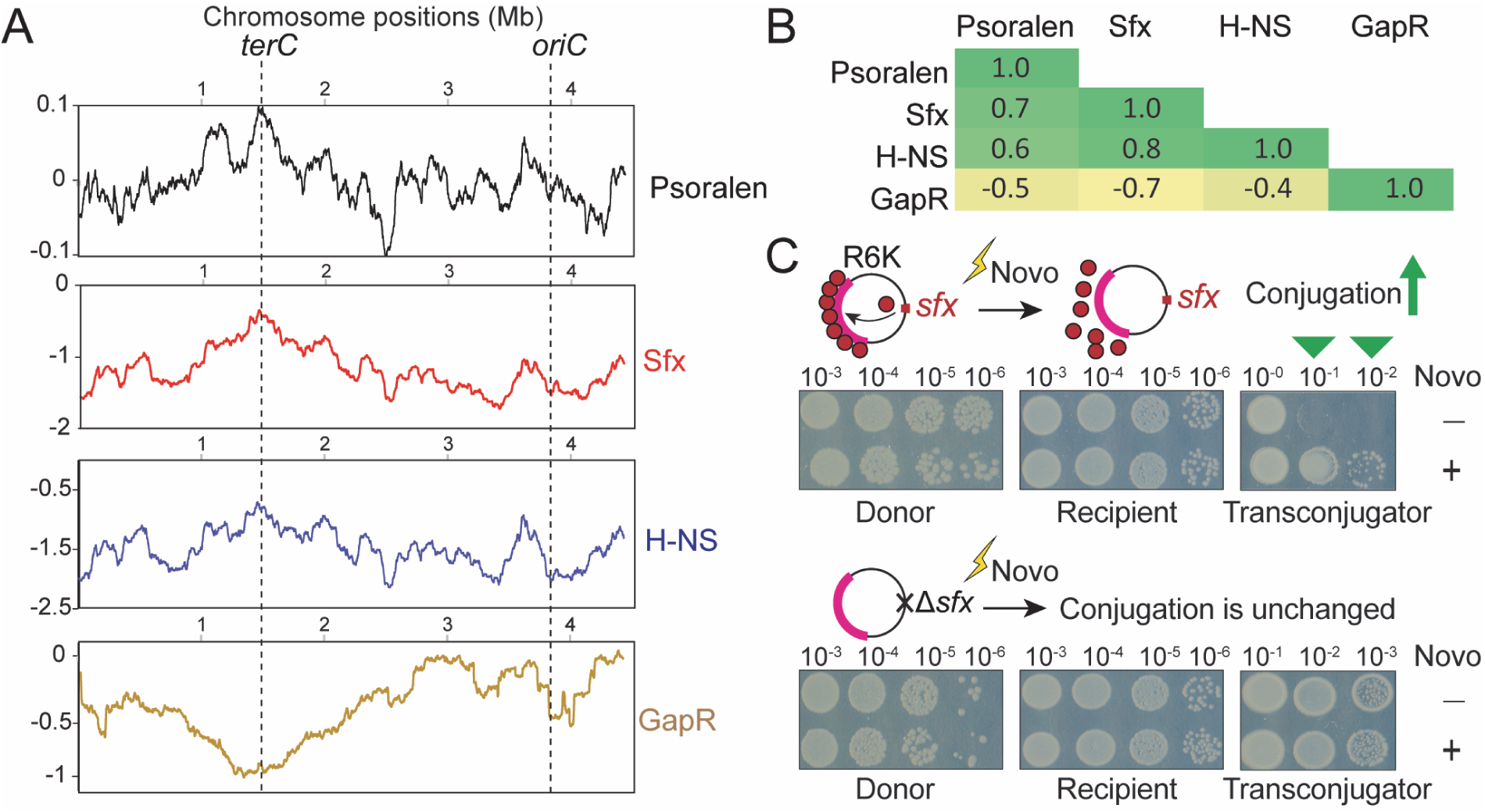
Sfx prefers negatively supercoiled DNA. (**A**) Correlation between DNA topology and Sfx/H-NS binding sites. The Y-axis values represent the moving average of 200 kbp windows. Psoralen plot is the log-transformed ratio of psoralen crosslinked to non-crosslinked fraction. Sfx/H-NS/GapR plots are the mean values of log2FE of two biological repeats. FE = ChIP/Input. Positions of the chromosome replication terminus (terC) and origin (oriC) are indicated. (**B**) Pearson correlation between the calculated moving average in panel **A**. (**C**) Effects of novobiocin (Novo) on R6K conjugation. Serial dilutions are indicated on the top of the plates. Novo treatment disrupts the Sfx-mediated *vir* operon (magenta arc) silencing. Representative results from three independent biological repeats are shown.

To determine whether Sfx activity in the cell depends on DNA topology, we used novobiocin, which inhibits DNA gyrase and negative supercoiling (85). We carried out conjugation assays with donors carrying the wild-type and Δ*sfx* R6K following a 1 h treatment with novobiocin. Our results demonstrate that the inhibition of DNA gyrase dramatically (>100-fold) increases the conjugation efficiency of wild-type R6K to match the level of R6KΔ*sfx* but has no effect on that of R6KΔ*sfx* (Fig. 3C). We conclude that the loss of negative supercoiling abolishes Sfx-mediated silencing (and likely targeted binding).

### Sfx mimics H-NS effects mediated by DNA and RNA interactions

Our findings that Sfx and H-NS share many binding sites on the chromosome are consistent with a previous report that Sfx can at least partially complement some phenotypes of the Δ*hns* strain, such as motility and growth defects (67). While H-NS has been extensively studied over the years, nothing is known about the molecular mechanism of Sfx. We thought that comparing the effects of Sfx and H-NS on genes where the mode of action of H-NS is known could shed light on that of Sfx. We first tested the ability of Sfx to silence the cryptic *bgl* operon, which enables *E. coli* to catabolize β-glucosides, the activity that can be assayed on MacConkey indicator agar supplemented with selected sugar. The *bgl* operon is one of the best-studied targets of H-NS, which binds to high-affinity DNA sites flanking the promoter and forms filaments that block both sense and antisense transcription (Fig. 4A) (12,58,86). H-NS inhibits RNAP binding to the promoter (86) and increases Rho-dependent termination by inducing RNAP pausing during elongation (58). The block to elongation requires the formation of bridged H-NS filaments (58) and is robustly stimulated by Hha and StpA, which form mixed filaments with H-NS (15). Unlike H-NS-only filaments, the StpA-only filaments efficiently blocked RNAP progression (15), a finding of particular interest given that Sfx is a closer paralog to StpA (67). Our results show that H-NS and Sfx have similar occupancy profiles on the *bglHGFB* genes (Fig. 4B) and, when expressed from their native promoters (as described in (67)), can restore the *bgl* operon silencing in the *Δhns E. coli* strain, as assayed on MacConkey salicin plates (Fig. 4B).

**Figure 4.**
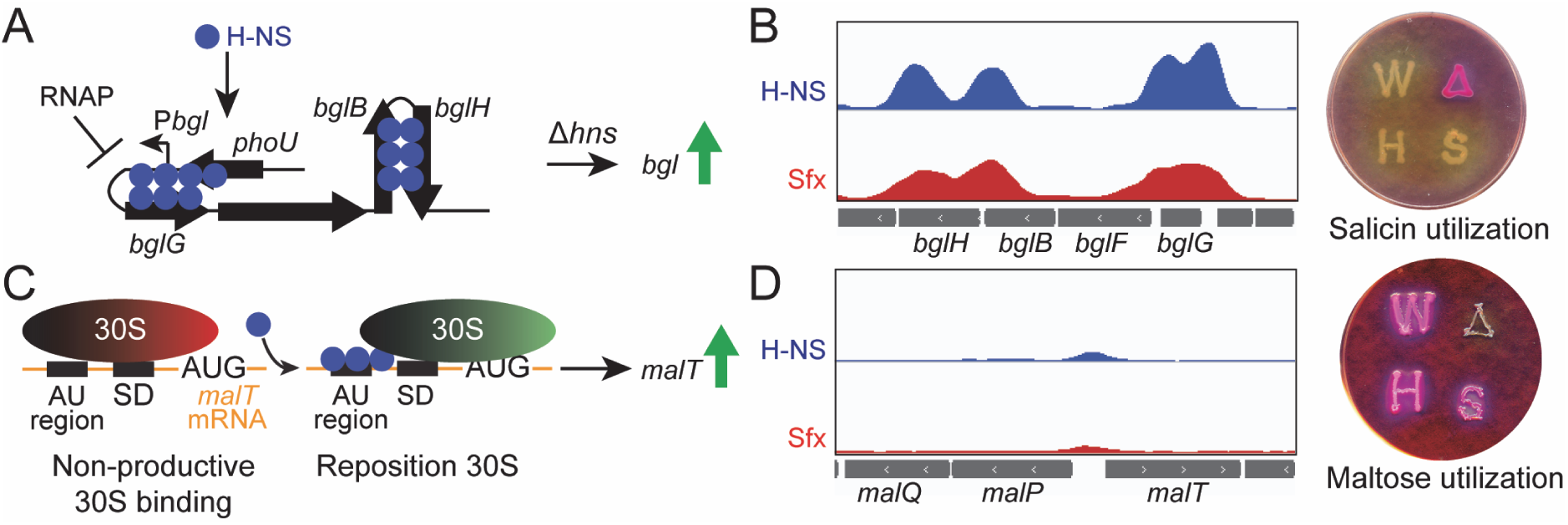
Sfx can complement the *hns* deletion phenotypes. (**A**) Mechanism of H-NS-mediated silencing of the *bgl* operon. Bridged H-NS filaments block transcription initiation and elongation. (**B, D**) Left, the tracks represent FE, which ranged from 0 to 7. Right, *in vivo* test results of sugar utilization using the MacConkey indicator agar. The *E. coli* Δ*hns* strain was transformed with plasmids expressing H-NS (H), Sfx (S), or an empty vector (Δ); the wild-type *E. coli* (W) was transformed with an empty vector. Single colonies were patched onto MacConkey plates supplemented with 0.6% salicin or 0.4% maltose and incubated for 20 hours at 30 °C; a representative plate (out of five biological replicates tested) is shown. (**C**) Mechanism of H-NS-mediated activation of the *mal* operon. The *malT* mRNA contains a suboptimal Shine-Dalgarno (SD) sequence. H-NS binds to the AU-rich region of the *malT* mRNA and repositions the 30S from a non-productive state to an active translation initiation state, thereby increasing the translation of MalT, the activator of the *mal* operon transcription.

Although most regulatory effects of H-NS are mediated through DNA interactions, H-NS and StpA bind RNA (87,88) and can control gene expression post-transcriptionally. For example, H-NS and StpA activate expression of the *E. coli* maltose regulon by increasing translation of *malT*, a gene that encodes the activator of the maltose regulon (89,90). This effect was shown to depend on H-NS-mediated 30S repositioning (Fig. 4C) on the *malT* mRNA (89). As expected, we observed that the *hns* deletion abolished maltose utilization (as assayed on MacConkey maltose plates; Fig. 4D). The ectopic expression of either *sfx* or *hns* restored maltose utilization, suggesting that Sfx may be able to bind RNA.

To identify potential RNA ligands of Sfx, we used RNA ChIP-seq, in which the samples were treated with DNase after crosslinking reversal to enrich for Sfx-bound RNAs, which were then sequenced. This analysis identified several enriched RNAs, such as *ssrA*, and we were able to confirm that Sfx interacts with the SsrA RNA *in vitro* (Supplementary Fig. S4). However, the physiological significance of this observation remains to be determined; for example, we did not observe enrichment in *malT* mRNA, even though our complementation results (Fig. 4D) indirectly support a model in which Sfx binds RNA. It is possible that Sfx pulls down more abundant RNAs; calculated from our RNA-seq data, the *ssrA* RNA is ∼268 times more abundant than *malT* RNA. Another reason is that most of our RNA ChIP-seq reads are rRNA, which reduces the chance of detecting *malT* RNA.

### Sfx is preferentially localized to the R6K vir operon

Our results demonstrate that Sfx has minimal effects on chromosomal gene expression (Fig. 1F) and apparently binds the chromosome less well than H-NS at most sites (Fig. 2C). However, the two proteins localize to similar AT-rich regions and recognize similar DNA motifs (Fig. 2E). This pattern is consistent with the ability of the ectopically-expressed Sfx to substitute for missing H-NS, as shown here (Fig. 4) and in the previous report (67), and reminiscent of other plasmid-encoded H-NS homologs (29).

Our analysis of Sfx and H-NS binding to R6K reveals a strikingly distinct pattern: Sfx forms two continuous multi-kb peaks along the *vir* operon, whereas H-NS is largely excluded (Fig. 5A), even though the *vir* operon is very AT-rich (Fig. 1A). Our ChIP-seq and RNA-seq data suggest that R6K is organized into three distinct domains (Fig. 5A). (1) The ARG domain, a transposition island that encodes three antibiotic resistance genes, is GC-rich and does not bind either H-NS or Sfx. (2) The PIR domain, which contains the highly transcribed *pir* gene, binds Sfx and H-NS equally well. Gene expression of the PIR domain is largely unaffected by Sfx; the only exception is *hyp2*, which is upregulated upon *sfx* deletion (Fig. 1F). This result, however, could be artefactual: in the R6K*Δsfx,* the *hyp2* gene is placed next to the *sfx* gene promoter (Fig. 5A, Dataset 2), located at the end of the *topBx* gene (which encodes a putative topoisomerase), and may become fortuitously activated. (3) The VIR domain, which includes the signature *vir* operon and other conjugation genes, is almost continuously covered by Sfx. Consistently, the conjugation function-related VIR domain genes are silenced by Sfx (Fig. 1AF).

**Figure 5.**
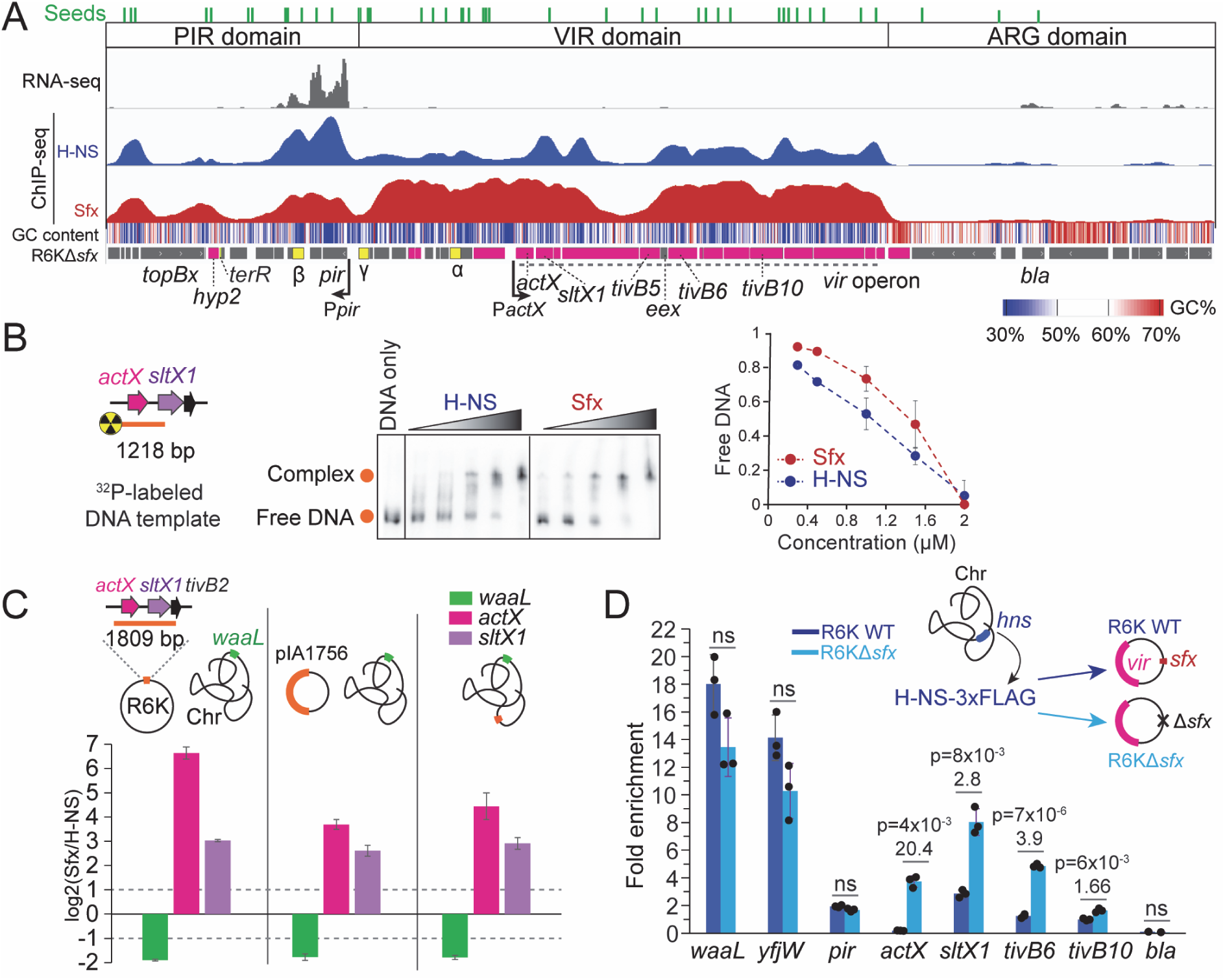
H-NS is largely excluded from the R6K *vir* operon. (**A**) RNA seq track of wild type R6K is shown along with H-NS and Sfx ChIP-seq tracks. H-NS and Sfx ChIP-seq tracks are in same scale for comparison. The predicted high-affinity binding sites (Seeds) are indicated as green bars on the top. The proposed three domains of R6K are indicated. The GC content, calculated in a 25-bp sliding window, is shown as a heatmap below the tracks. Significantly upregulated genes by *sfx* deletion are highlighted in magenta in the R6KΔ*sfx* map. The promoters of *pir* (P*pir*) and the *vir* operon (P*actX*) are indicated. The positions of three replication origins are shown in yellow, and the replication terminus (*terR*) in lime yellow. (**B**) Interactions of Sfx and H-NS with the *actX-sltX1* fragment were tested by EMSA. Protein concentrations ranged from 0.3 to 2 μM; the radioactively labeled DNA was fixed at 35 nM. The proportion of free DNA is plotted against protein concentrations; error bars are the SD (n = 3). (**C**) Moving *actX-sltX1-tivB2* fragment from R6K into plasmid pIA1756 (Supplementary Fig. S7) and chromosome. ChIP-qPCR results show the enrichment of selected fragments. FE (ChIP/Input) was first calculated using Cq from qPCR, and the log2(Sfx FE/H-NS FE) is shown as bar graph. Error bars are the SD (n = 3). (**D**) ChIP-qPCR quantifies the FE of H-NS on different genes. The FE, calculated for both wild type and Δ*sfx* R6K, are shown as bar graph. Error bars are SD (n = 3). A two-tailed t-test assuming unequal variances was used to calculate p-values. Fold change (FC) was calculated as FE (R6KΔ*sfx*) / FE (WT R6K). Values for genes with p-value < 0.05 and FC > 1.5 are indicated at the top of bars, and “ns” is indicated otherwise.

The Sfx VIR footprint is demarcated by the main origin of R6K replication, γ *ori*, and the highly transcribed *pir* gene on one side, and by a “red” wall (> 65% GC, with a potential to form G-quadruplets) on the other side (Fig. 5A), features that could act as barriers to the Sfx filament extension. Notably, a gap in Sfx and H-NS occupancy occurs near the center of the VIR peak (Fig. 5A), in a region characterized by the typical AT-rich content of the *vir* operon (Fig. 1A). At least two mechanisms could explain this discontinuity in the Sfx filament. First, each VIR domain peak contains many potential high-affinity Sfx-binding motifs (Seeds in Fig. 5A), whereas the valley only has two (Fig. 5A). If Sfx requires several Seeds to form stable filaments, it could be outcompeted by other proteins in Seed-poor regions; for example, H-NS is excluded from CHIN loops by HU (91). While Sfx appears to exclude HU on the chromosome (Supplementary Fig. S5), many other DNA-binding proteins could perform a similar exclusionary role (92) to block Sfx binding. Second, we speculate that transertion, a process in which transcription, translation, and insertion of a protein into the membrane are closely coupled (93), anchors the VIR domain DNA to the cytoplasmic membrane, effectively partitioning this region into two Sfx-interacting VIR subdomains. The transertion-dependent bridges between the DNA and the membrane are known to play key roles in the domain organization of the chromosomal DNA (94) and of model plasmids, such as pBR322 (95). Two genes located in/near the valley (Fig. 5A), *tivB5* and *eex*, are expected to undergo transertion. The *tivB5* gene encodes a putative adhesin that binds to specific receptors on the surface of the target cell (96) and must traverse both cellular membranes *en route* to its location at the tip of the conjugative pilus (97). The *eex* gene encodes a putative entry exclusion factor, a membrane lipoprotein that impedes plasmid transfer to the recipient bacterium that already carries a similar conjugative plasmid to protect the cell from lethal zygosis, cell death induced by many rounds of conjugation that damage the cell wall (98,99). Studies of F and other Inc-type plasmids revealed that TivB5 and Eex act through specific contacts to the donor’s TraG/VirB6/TivB6 protein (97,100,101). Interestingly, Sfx has only a modest effect on *tivB5* and *tivB6* expression and no effect on *eex* (Fig. 1F), suggesting that these genes are at least partially independent of Sfx control.

Our analysis of the chromosomal occupancy patterns of Sfx and H-NS shows that they recognize similar AT-rich sequences and that H-NS binds to the chromosome better than Sfx (Fig. 2C-E); we did not observe a single instance of a higher Sfx peak on the chromosome. The reversal of this pattern in the R6K VIR domain (and to a lesser extent in the PIR domain) (Fig. 5) implies the existence of mechanisms that mediate preferential recruitment of Sfx or selective exclusion of H-NS.

### VIR domain preferentially binds Sfx on supercoiled DNA

To explain how Sfx is selectively targeted to the VIR domain, we examined several possibilities. *First*, Sfx may be preferentially recruited to R6K *in cis*. However, the published data (67) and our results demonstrate that Sfx expressed *in trans* from its cognate promoter on a low copy number plasmid (Fig. 2B and Supplementary Fig. S3B) silences R6KΔ*sfx* conjugation refute this explanation.

*Second*, the R6K DNA may contain modifications that inhibit H-NS, but not Sfx binding. This explanation appears farfetched since (1) R6K does not encode a putative DNA modification enzyme and (2) the modification would be restricted to the VIR domain. Nonetheless, we compared H-NS binding to plasmid DNA extracted from cells to PCR-amplified DNA (Supplementary Fig. S6) and did not observe any differences, excluding DNA modification as the mechanism of H-NS exclusion.

*Third*, it is possible that H-NS prefers to bind the chromosome and is present in insufficient quantities to cover R6K. This explanation appeared unlikely because H-NS binds better than Sfx to some R6K regions, for example, the *pir* gene and the β *ori* (Fig. 5A), and R6K increases the total DNA content of the cell only by 14%, an increase that may be offset by autoregulation of H-NS (71).

*Fourth*, Sfx may bind the VIR domain with higher affinity than H-NS. Using a linear 1218 bp DNA fragment encompassing the *actX-sltX1* region, which is strongly associated with Sfx (Fig. 5AB), we found that Sfx and H-NS bind equally well (Fig. 5B). To test whether Sfx binding would be selectively favored on supercoiled DNA, we inserted the *actX-virB1* fragment into an unrelated 4 kb plasmid (pIA1756; Supplementary Fig. S7A) and assayed the H-NS and Sfx binding with ChIP-qPCR (the positions of probes are shown in Supplementary Fig. S7B). Our results reveal that Sfx binds the chromosomal *waaL* gene less well than H-NS (Fig. 5C), in line with the ChIP-seq observations (Fig. 2D). In contrast, Sfx preferentially binds to the *actX* and *sltX1* genes on the plasmid (Fig. 5C). We next moved a fragment containing *actX* and *sltX1* genes to the chromosome, choosing the CP4-57 prophage region to which H-NS binds well, but Sfx does not (Fig. 2D); again, we found that Sfx binds *actX* and *sltX1* genes better than H-NS (Fig. 5C). These results demonstrate that Sfx preferentially binds to the VIR domain segment located on the negatively supercoiled DNA (either the chromosome or a plasmid), but not on the linear DNA.

We hypothesize that Sfx recognizes a specific structure that forms in the topologically constrained VIR domain. Recognizing that probing the VIR domain DNA structure in the cell would be challenging, we carried out two types of experiments that lend support to this hypothesis. *First*, we showed that the inhibition of DNA gyrase by novobiocin abolishes conjugation silencing by Sfx (Fig. 3C). *Second*, it has been shown that melting and the transition from B- to L-form DNA are sensitive to GC content (102,103). The 4 kb plasmid (pIA1756, 44% GC content) that contains the *actX-sltX1* fragment and selectively binds Sfx (Fig. 5C) displayed highly compact topologies in AFM images, as did a similarly sized control plasmid (pBR322, 54% GC) (Supplementary Fig. S7C), whereas a 5.8 kb plasmid pDM_N1_400 (54% GC; the pBR322 backbone) shows a relaxed topology (Supplementary Fig. S7C). Ongoing single-molecule studies of larger segments of the VIR domain may reveal distinct topology.

Does this unusual structure block H-NS binding to the R6K VIR domain? To explore this possibility, we compared H-NS binding to selected genes on the wild-type R6K and R6KΔ*sfx* using ChIP-qPCR. As controls, we used the chromosomal *waaL* and *yfjW* genes, which are strongly bound by H-NS (Fig. 2D). We found that H-NS binding to the chromosome was not significantly affected by the *sfx* deletion on R6K, as could be expected. We also did not observe changes in H-NS association with the PIR and ARG domains of R6K (Fig. 5D), using the *pir* and *bla* genes as probes, respectively. In sharp contrast, H-NS binding to the *vir* operon was notably increased upon the deletion of *sfx* from R6K: the H-NS ChIP signal enrichment in *actX* increased 20 times, and 3 – 4 times for *sltX1* and *tivB6* genes (Fig. 5D). We conclude that the selective H-NS exclusion from the R6K VIR domain is mediated by Sfx/DNA complexes, which assemble on the negatively supercoiled DNA.

### Phase separation promotes H-NS exclusion from R6K

Sfx is predicted to be folded similarly to H-NS and to contain dimerization, oligomerization, and DNA-binding modules, but their length and orientation may differ (Fig. 6A). R6K Sfx and ∼73% of Sfx homologs from conjugative IncX plasmids have predicted IDRs, whereas *E. coli* H-NS does not (Figs. 1B, 6A). IDRs are often involved in phase separation (104), a phenomenon that creates membrane-less organelles in cells (105). We wondered whether phase separation can be utilized to exclude H-NS from Sfx-R6K. We first used a pelleting assay to examine whether Sfx can form phase-separated condensates. We found that purified Sfx can be efficiently pelleted with R6K or total RNA (Fig. 6B), whereas H-NS, less efficiently, is pelleted with R6K in the absence (Fig. 6B, left), but not in the presence of Sfx (Fig. 6B, right), supporting the idea that Sfx outcompetes H-NS for binding to R6K. These results can be explained by macromolecular aggregation or phase separation.

**Figure 6.**
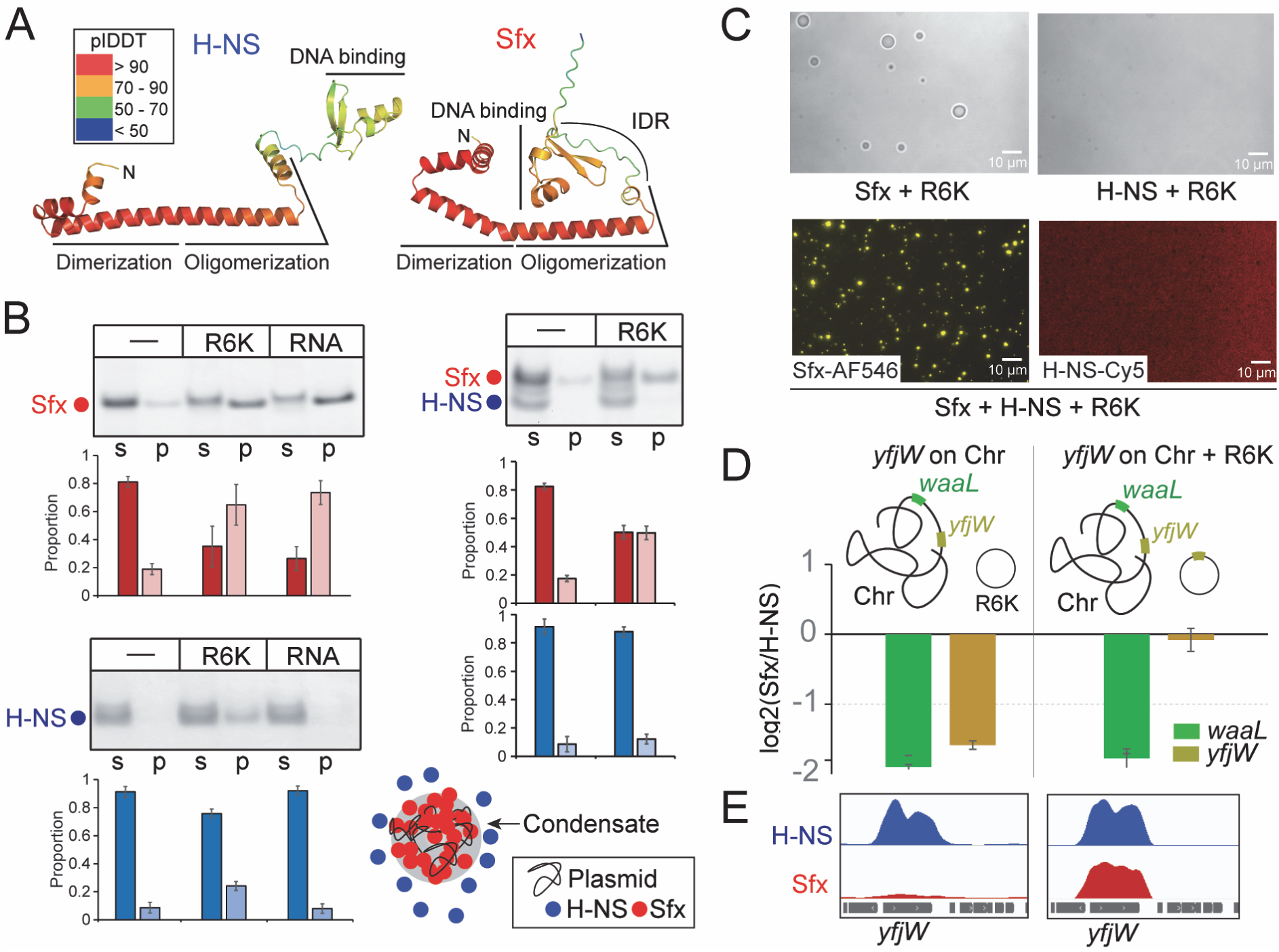
Phase separation of Sfx and R6K. (**A**) Structural models of Sfx and H-NS were predicted by AlphaFold3. (**B**) Pelleting assays of Sfx and H-NS with R6K or RNA. (Left), Sfx and H-NS were tested separately. The RNA is a total RNA extracted from the exponential phase of *E. coli* cells. Each protein (at 2 μM) was pelleted with 5 ng/μl R6K or 30 ng/μl total RNA. s, supernatant. p, pellet. (Right) Sfx and H-NS were mixed. Bar graphs (Sfx, red; H-NS, blue with the darker shade corresponding to the soluble fraction) were generated from three biological repeats; error bars are the SD. (**C**) Microscopy of Sfx phase separation with R6K. In the presence of 3% PEG8000, 2 μM unlabeled proteins were mixed with 0.5 μM fluorescently labeled proteins and 5 ng/μl R6K. (Top), Sfx and H-NS are tested individually. Bright field views are shown. (Bottom) Sfx and H-NS are mixed. Fluorescence views were obtained for AF546-labeled Sfx and Cy5-labeled H-NS. Scale bar, 10 μm. (**D**) ChIP-qPCR results of the *yfjW* gene enrichment after moving it into R6K. FE (ChIP/Input) was first calculated using Cq from qPCR, and then the log2(Sfx FE/H-NS FE) is shown as bar graph. Error bars are the SD (n = 3). (**E**) ChIP-seq tracks of the *yfjW* gene. Left, ChIP-seq with *yfjW* on the chromosome. Right, ChIP-seq with *yfjW* on the chromosome and R6K. Peaks represent FE, and the same scale is used for both H-NS and Sfx tracks within each scenario.

To evaluate the possibility of phase separation, we utilized microscopy. From bright-field microscopy, we find that Sfx forms condensates with R6K and RNAs, while in the same conditions, HNS does not (Fig. 6C, top and Supplementary Fig. S8A). Next, we asked if H-NS can enter the Sfx-R6K condensate. Pelleting assays show that when Sfx, H-NS, and R6K are present together, Sfx is pelleted while H-NS remains mostly in the supernatant (Fig. 6B); 300 mM KCl, but not 5% 1,6-Hexanediol, disrupts the formation of Sfx-R6K pellet (Supplementary Fig. S8B), indicating that the assembly is mainly driven by electrostatic interactions. The fluorescence microscopy analysis of samples containing AF546-labeled Sfx, Cy5-labeled H-NS, and R6K shows that Sfx is concentrated inside droplets, whereas H-NS is evenly distributed (Fig. 6C, bottom). However, H-NS does not appear to be selectively excluded from Sfx-R6K condensates (Fig. 6C), consistent with the ChIP-seq data showing that H-NS weakly binds to the VIR domain (Fig. 5A).

These *in vitro* assays suggest that phase separation could assist Sfx in safeguarding the R6K VIR domain from nonproductive (in terms of gene expression silencing) association with H-NS indirectly, by sequestering Sfx within a microcompartment formed by up to 16 copies of R6K. If this compartmentalization occurs, Sfx should possess a competitive binding advantage for any gene located on the R6K plasmid. To test this, we used the *yfjW* gene from the CP4-57 prophage, which exhibits the highest chromosomal H-NS occupancy and minimal Sfx binding, indicating that Sfx binding to *yfjW* is outcompeted by H-NS (Fig. 2D). We postulated that moving the *yfjW* gene to R6K should favor binding of Sfx. We used ChIP-qPCR to compare two scenarios (1) *yfjW* on the chromosome only, and (2) *yfjW* present on both the chromosome and within the R6K ARG domain (Fig. 6D). We found that while the relative binding of H-NS and Sfx to the *waaL* gene remained constant across both scenarios, Sfx recruitment to *yfjW* shifted dramatically. Sfx crosslinked poorly to *yfjW* in the first scenario but bound *yfjW* as well as H-NS in the second scenario (Fig. 6D). This occupancy trend was confirmed by ChIP-seq analysis, demonstrating that Sfx efficiently binds *yfjW* specifically on R6K (Fig. 6E). These findings support our model that phase separation provides Sfx with a localized competitive advantage.

## DISCUSSION

Many bacteria utilize two (or more) paralogous histone-like proteins to silence horizontally transferred genes (HTG). Examples include H-NS and StpA in *E. coli*, MvaT and MvaU in *Pseudomonas aeruginosa*, and Lsr2 and LsrL in Actinobacteria (11). The secondary protein (StpA, MvaU, LsrL) may appear partially redundant, having fewer targets than the major silencer (68,106–110). In addition, hundreds of H-NS homologs have been identified on plasmids (111). It remains unclear how these silencers coordinate or if the “redundant” paralogs possess specialized functions.

The relationship between the transcription elongation factors NusG and RfaH, which modulate HTG silencing, provides a model for such specialization. While the essential NusG works with Rho and H-NS to silence broad sets of HTG (12,16,112), its paralog RfaH is recruited to the elongating RNAP at specific DNA sequences in a small subset of foreign operons (17,20) to override silencing (19,22,23). Here, we demonstrate that the F plasmid conjugal transfer (*tra*) operon is antagonistically regulated by RfaH and Rho (Supplementary Fig. S1).

Evidence suggests histone-like proteins may be specialized; for instance, despite sharing DNA-binding preferences and transcriptional effects with H-NS (113), StpA functions as a potent RNA chaperone, an activity that relies on weak and transient interactions (88,114,115), in contrast with rigid DNA-bridging filaments that hinder RNAP elongation by H-NS and StpA (15,58,116).

Here, we examined an interplay between H-NS and its homolog, Sfx, which is encoded on R6K plasmid and serves to suppress its conjugation. A recent report highlights a striking functional asymmetry: while Sfx can replace chromosomal H-NS functions, H-NS is incapable of silencing R6K transfer (67). Our results show that Sfx acts synergistically with Rho to block the expression of the R6K conjugal transfer *vir* operon during RNA chain elongation but does not inhibit initiation at the major *vir* operon promoter (Fig. 1DE). This finding is surprising because Sfx-like proteins have been shown to act at the promoter (54,55). Our ChIP-seq analysis reveals that, on the chromosome, Sfx and H-NS binding sites largely overlap; both proteins recognize similar DNA motifs (Fig. 2CE) and have an apparent preference for the negatively supercoiled DNA (Fig. 3). However, Sfx occupancy is lower at most sites (Fig. 2CD) and, consistently, Sfx deletion has little effect on the chromosomal gene expression (Fig. 1F). On R6K, this pattern is drastically different: Sfx is strongly enriched across the entire VIR domain, which includes the *vir* operon, whereas H-NS is largely excluded (Fig. 5A). Curiously, Sfx and H-NS occupancies in the PIR domain are comparable, which could arise from the presence of the γ origin and replication terminus *terR* (117,118) (Fig. 5A). This idea is based on our observation that similar Sfx and H-NS ChIP peaks are often found around the replication origin and terminus on the chromosome (Fig. 2C). As discussed in the Results section, the true protein occupancies cannot be inferred from ChIP peak analyses; we thus focus on relative differences between H-NS and Sfx binding to DNA.

### Hypothetical mechanisms of preferential Sfx targeting to R6K

Our data support two mutually reinforcing mechanisms that explain the observed selectivity of Sfx (Fig. 7). First, we hypothesize that the *vir* DNA forms a unique architecture recognized by Sfx as a high-affinity loading site, followed by the spread of Sfx filaments along the DNA, a shared behavior of H-NS homologs (11,74,75). We demonstrate that Sfx cellular activity depends entirely on negative DNA supercoiling (Fig. 3). Single-molecule (119) and high-resolution chromosome conformation capture (91) experiments demonstrate that H-NS pins plectonemes *in vitro* and *in vivo*, bridging chromosomal hairpin (CHIN) stems (91). Our ChIP-seq data reveal a distinct binding pattern: H-NS occupancy tracks with CHIN stems and loops (peaks and valleys), whereas Sfx forms uniform, wider peaks across the entire hairpin (Fig. 2G, Supplementary Fig. S5).

**Figure 7.**
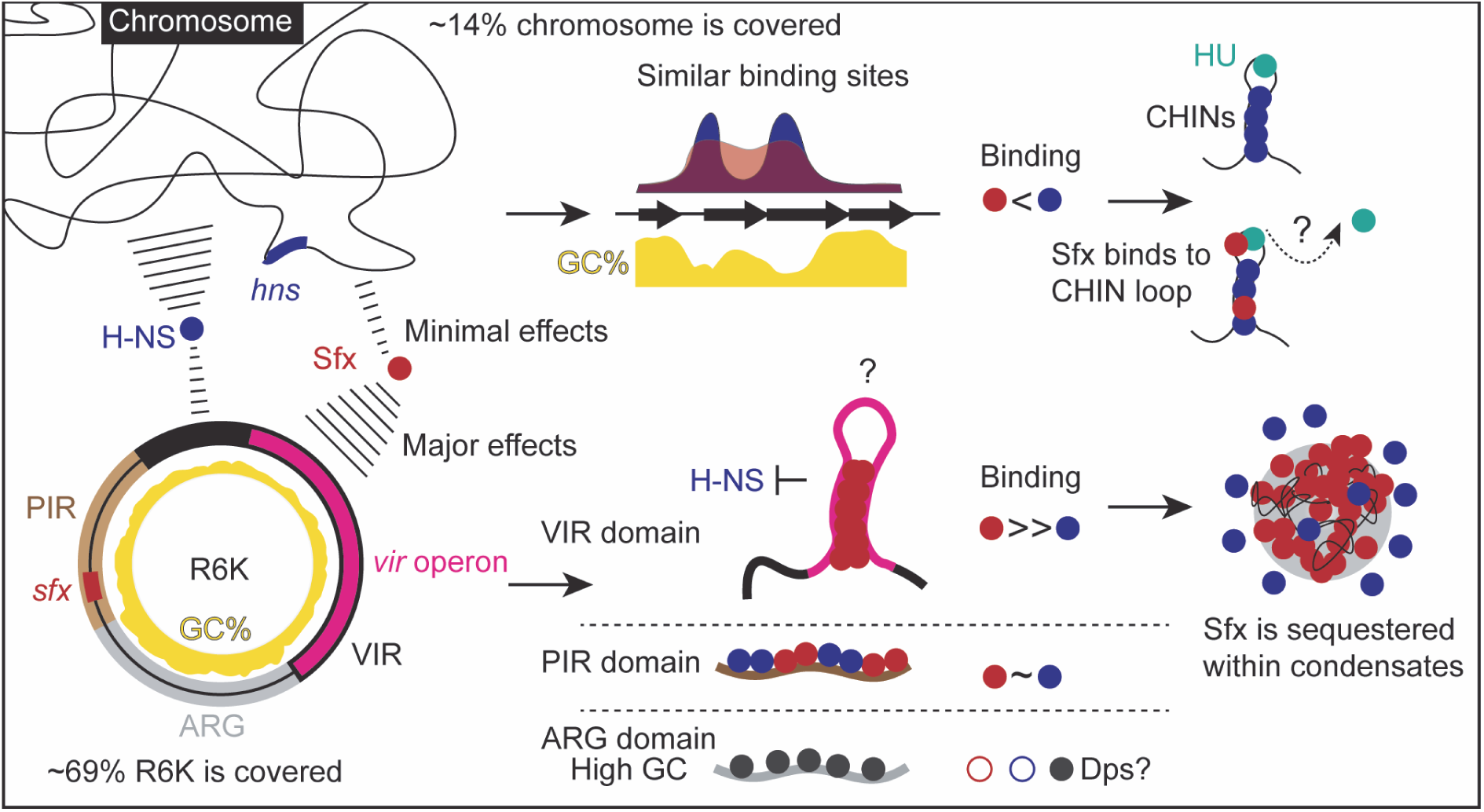
The proposed model of selective targeting of Sfx to the R6K conjugation functions. While Sfx strongly silences the R6K *vir* operon, H-NS exerts minimal influence, even when Sfx is absent. Although H-NS and Sfx occupy similar chromosomal sites, Sfx uniquely extends into CHIN loops to potentially displace HU. Our analysis of the R6K plasmid reveals three distinct domains that differ in their binding to H-NS and Sfx. The PIR domain binds both proteins, while the ARG domain binds neither, potentially using Dps to fill the void. Notably, Sfx far outnumbers H-NS in the VIR domain. We propose that H-NS is excluded from the VIR domain through two primary drivers: the inherent DNA structure of the *vir* operon, which promotes the formation of stable Sfx filaments, and the phase separation of Sfx-R6K complexes. See text for additional details.

Single-molecule experiments on synthetic DNAs show that AT-rich sequences pin plectonemes at specific locations (120) to determine the overall DNA structure. Modeling indicated that DNA deformability, which can be calculated from dinucleotide steps, rather than overall AT content, underlies this behavior. The *vir* operon has a high AT content and many deformable dinucleotide steps, favoring conformational flexibility, and *vir* DNA may form compact structures (Supplementary Fig. S7C). Future experiments will determine the location and structure of the high-affinity Sfx binding site(s) that we propose mediate its initial loading on R6K (Fig. 5A).

Second, we hypothesize that phase separation ensures that Sfx is sequestered within R6K condensates, promoting initial Sfx binding and spreading along the VIR domain and preventing Sfx “escape” to the chromosome, which has an excess of Sfx binding motifs. The Sfx sequestration would be expected to favor its binding to R6K. Indeed, Sfx binds better to the *actX* gene on R6K as compared to the same region placed on the chromosome or an unrelated plasmid (Fig. 5C). Strikingly, Sfx gains the ability to bind the *yfjW* gene, which shows no Sfx occupancy on the chromosome (Fig. 2D), once *yfjW* is moved to R6K (Fig. 6DE). This sequestration will further bolster the competitive advantage of Sfx over H-NS, indirectly excluding H-NS from the preformed Sfx/DNA complexes. Pelleting and microscopy assays show that Sfx forms condensates with RNA and DNA (Fig. 6BC and Supplementary Fig. S8). While H-NS can also phase separate at high concentrations, as shown by others (121), Sfx does so more readily. These condensates are likely mediated by the IDR found in Sfx but absent in H-NS (Fig. 6A), a possibility that we will explore in future experiments. While (to our knowledge) phase separation has not been reported for the Sfx-homolog StpA, it also possesses a large degree of intrinsic disorder, and its weak and dynamic contacts with RNA are thought to underpin its ability to act as an RNA chaperone (114).

### Uninterrupted nucleoprotein filaments may be required to silence R6K conjugation

We posit that the efficient silencing of conjugation requires 100% occupancy of the *vir* operon by Sfx-only or mixed Sfx/H-NS filaments (Fig. 5A). The presence of mixed filaments is suggested by the H-NS ChIP signals observed within the continuous Sfx footprint. However, the relative levels of H-NS in these hypothetical mixed filaments may be lower than appears from the ChIP-seq peaks (see above about the discussion of ChIP efficiency).

Does the cellular concentration of Sfx suffice to fully cover the *vir* operon? Based on our ChIP-seq data, we estimate the potential binding areas for H-NS and Sfx as ∼0.6 Mb on the chromosome (primarily H-NS bound) and ∼0.4 Mb on R6K (the aggregate of 16 copies), respectively (Supplementary Fig. S3D). Our findings align with previous reports that *E. coli* H-NS covers 10% – 15% (0.46–0.7 Mb) of the chromosome (122). Based on *in vitro* measurements, one H-NS dimer is expected to cover ∼15 bp of DNA (123). Assuming that Sfx makes similar contacts to DNA, ∼27,000 Sfx dimers would be needed to saturate 0.4 Mb of R6K DNA. While direct absolute measurements for Sfx are unavailable, quantitative proteomics shows that H-NS is present at 22,000 copies in the stationary phase *E. coli* (124). Since Sfx is present in at least a 3.5-fold excess (Supplementary Fig. S3D), the total count of Sfx dimers exceeds 38,000 per cell. This abundance is more than sufficient to provide complete coverage of R6K, with a surplus of dimers available for binding to the chromosome.

Our observations that Sfx promotes early transcription termination *in vivo* (Fig. 1E) suggest that it forms bridged filaments on the DNA, since only bridged filaments can efficiently block the elongating RNAP (58). Our ChIP-seq data show that, compared to H-NS, Sfx forms wider peaks along the same DNA regions (Fig. 2FG, Supplementary Fig. S5). It is plausible that Sfx forms longer, more stable filaments, which may displace other chromatin-associated proteins from the DNA; we found that Sfx binds to CHIN loops thought to be occupied by HU (91).

Differences in filament structure could explain why H-NS does not inhibit conjugation even though it binds the *vir* operon in the absence of Sfx (Fig. 5D): a break in the filament may enable recruitment of RNAP or accessory proteins and destabilize the filament further. A precedent for this exists in *Shigella* VirB, which binds to and spreads on DNA, removing negative supercoils and abrogating H-NS silencing (79). Curiously, VirB is a member of the ParB family of DNA partitioning proteins, and R6K encodes a distant homolog of ParB.

While H-NS forms bridged filaments under some conditions, this property is promoted by Hha (H-NS-associated protein), a small (∼8 kDa) co-regulator that forms mixed filaments with H-NS (15,125). Hha enhances the ability of H-NS to arrest the elongating RNAP (15,23) and increases the Sfx-mediated silencing of R6K conjugation (67). To investigate Hha’s influence on Sfx-R6K binding, we performed ChIP-seq in an *hha* deletion strain. We observed that while the specific binding positions of Sfx on both R6K and the chromosome remained unchanged (Supplementary Fig. S9A), the peak heights on R6K were noticeably reduced in the absence of Hha (Supplementary Fig. S9B). Peak heights of Sfx on the chromosome increased in the absence of Hha (Supplementary Fig. S9C), possibly arising from the decreased H-NS binding. These findings suggest that Hha and Sfx may co-assemble into mixed bridged filaments to achieve more potent transcriptional silencing. Consistently, Hha is pelleted together with Sfx (Supplementary Fig. S9D) at a 1:1 ratio, as observed with the H-NS/Hha complex (125). Sfx paralog StpA also forms bridged filaments that efficiently block the RNAP progression, at least *in vitro* (15). Modeling by AlphaFold 3 (126) shows that StpA and Sfx form similar oligomeric structures, while H-NS does not (Supplementary Fig. S10).

### Structural organization and regulation of R6K

R6K consists of three modules: a GC-rich ARG domain flanked by the AT-rich PIR and VIR domains (Fig. 7). The ARG domain does not bind either H-NS or Sfx (Fig. 5A); we hypothesize that this region is bound by Dps, the most abundant nucleoid-associated protein in the stationary phase (127) whose occupancy on the chromosomal DNA (128) is inversely correlated with that of H-NS (Supplementary Fig. S11). The PIR domain binds both Sfx and H-NS, whereas the VIR domain is almost continuously covered by Sfx and largely lacks H-NS (Fig. 5A). Our ChIP-seq data suggest that the VIR domain forms a plectoneme; Sfx occupies and likely bridges the plectoneme stems but does not bind to the loop region (Fig. 5A), which may be anchored to the membrane through transertion (see *Sfx is preferentially localized to R6K vir operon*).

In our model (Fig. 7), Sfx represses R6K conjugation directly, by roadblocking the transcribing RNAP, and indirectly, by excluding H-NS from the transfer operon. The second mechanism may be unexpectedly critical, given that H-NS, the major xenogeneic silencer known to target numerous foreign genes, is unable to silence R6K (67) for yet-unknown reasons. While many other DNA-binding proteins have been implicated in the silencing of AT-rich xenogenes in *E. coli* (129), none appears able to substitute for Sfx, whose loss leads to ∼1,000-fold increase in R6K conjugation (67).

The multimodal silencing may be crucial for R6K, and in turn the host cell, because the loss of silencing filaments even on a single plasmid copy would lead to leaky expression of the T4SS genes, rendering the host vulnerable to pilus-specific phages. Accordingly, all 16 copies of R6K must be coordinately controlled by Sfx to ensure that conjugation is fully silenced. We propose that, in addition to conventional silencing of RNA synthesis, Sfx brings all R6K copies together by cross-bridging different plasmid molecules and promoting the formation of membrane-less cellular organelles. This multiprong strategy renders R6K self-sufficient and ensures its maintenance in diverse hosts, from *E. coli* to *Pseudomonas* (130), whose resident chromosomal silencers may fail to efficiently repress the conjugal transfer genes, as is the case with H-NS.

Plasmids employ sophisticated ’stealth’ mechanisms to mitigate host fitness costs. The H-NS homolog Sfh (pSfR27) serves as a classic molecular backup for H-NS, filling the chromosomal gaps in H-NS coverage caused by its repositioning to pSfR27 (28,29), and R6K Sfx can also act as a molecular mimic of H-NS (Fig. 4). However, our findings reveal that Sfx utilizes a superior stealth strategy to combine near-zero impact on host gene expression with very potent silencing of the plasmid genes. Sfx clusters with R6K to silence the plasmid genes and reduce its off-target binding to the chromosome, while preventing H-NS dilution from the host genome and non-productive interactions with R6K. This precision targeting avoids the pitfalls of molecular mimicry, effectively neutralizing the fitness costs of plasmid carriage.

## ACKNOWLEDGEMENTS

We are indebted to Steve Bell, Lydia Freddolino, Natacha Ruiz, and Will Navarre for many insightful discussions, and Rachel Samson for help with fluorescence microscopy. We also gratefully acknowledge gifts of plasmids (Will Navarre) and strains (Lydia Freddolino, Natacha Ruiz, and Joe Wade) that were used in this study. We acknowledge Killian Kirkpatrick for assistance with AFM.

## Author contributions

B.W. conducted the investigation, developed methodology, analyzed sequencing and other data, and prepared figures. R.G., N.B., B.K., N.M., C.J., D.D., and I.A. conducted the investigation and analyzed data. I.A and L.F. supervised the study and acquired funding. B.W. and I.A. wrote the manuscript. All authors edited the manuscript.

## SUPPLEMENTARY DATA

Supplementary data are available at NAR online.

Dataset 1: (A) Sfx distribution analysis in IncX group plasmids. (B) Differential gene expression analysis result of the RNA-seq data.

Dataset 2: GenBank format files of plasmid maps used in this study. The description of each plasmid is included in Supplementary Table S1.

## DATA AVAILABILITY

All plasmids and strains are available upon request. The NGS data generated in this study are deposited at the Gene Expression Omnibus with accession numbers GSE319657 for RNA-seq data and GSE319658 for ChIP-seq data.

Psoralen crosslinking microarray data (GSE77687) (82), GapR ChIP-seq (GSE152880) (83), HupB ChIP-seq (GSE181767) (131), and Dps ChIP-seq (GSE293552) (128) are available from NCBI.

## REFERENCES

1. Yang, L., Mai, G., Hu, Z., Zhou, H., Dai, L., Deng, Z. and Ma, Y. (2023) Global transmission of broad-host-range plasmids derived from the human gut microbiome. Nucleic Acids Res, 51, 8005–8019.

2. Larsson, D.G.J. and Flach, C.F. (2022) Antibiotic resistance in the environment. Nat Rev Microbiol, 20, 257–269.

3. Rodriguez-Beltran, J., DelaFuente, J., Leon-Sampedro, R., MacLean, R.C. and San Millan, A. (2021) Beyond horizontal gene transfer: the role of plasmids in bacterial evolution. Nat Rev Microbiol, 19, 347–359.

4. Paillard, P., Rouger, Q., Thomet, M. and Mace, K. (2025) Type IV secretion systems: from structures to mechanisms. EMBO J, 44, 6304–6319.

5. Thomas, C.M., Thomson, N.R., Cerdeno-Tarraga, A.M., Brown, C.J., Top, E.M. and Frost, L.S. (2017) Annotation of plasmid genes. Plasmid, 91, 61–67.

6. Raivio, T.L. and Silhavy, T.J. (2001) Periplasmic stress and ECF sigma factors. Annu Rev Microbiol, 55, 591–624.

7. San Millan, A. and MacLean, R.C. (2017) Fitness Costs of Plasmids: a Limit to Plasmid Transmission. Microbiol Spectr, 5.

8. Quinones-Olvera, N., Owen, S.V., McCully, L.M., Marin, M.G., Rand, E.A., Fan, A.C., Martins Dosumu, O.J., Paul, K., Sanchez Castano, C.E., Petherbridge, R. et al. (2024) Diverse and abundant phages exploit conjugative plasmids. Nat Commun, 15, 3197.

9. Koraimann, G. (2018) Spread and Persistence of Virulence and Antibiotic Resistance Genes: A Ride on the F Plasmid Conjugation Module. EcoSal Plus, 8.

10. Ray-Soni, A., Bellecourt, M.J. and Landick, R. (2016) Mechanisms of Bacterial Transcription Termination: All Good Things Must End. Annu Rev Biochem, 85, 319–347.

11. Singh, K., Milstein, J.N. and Navarre, W.W. (2016) Xenogeneic Silencing and Its Impact on Bacterial Genomes. Annu Rev Microbiol, 70, 199–213.

12. Peters, J.M., Mooney, R.A., Grass, J.A., Jessen, E.D., Tran, F. and Landick, R. (2012) Rho and NusG suppress pervasive antisense transcription in Escherichia coli. Genes Dev, 26, 2621–2633.

13. Lawson, M.R., Ma, W., Bellecourt, M.J., Artsimovitch, I., Martin, A., Landick, R., Schulten, K. and Berger, J.M. (2018) Mechanism for the Regulated Control of Bacterial Transcription Termination by a Universal Adaptor Protein. Mol Cell, 71, 911–922 e914.

14. Shen, B.A. and Landick, R. (2019) Transcription of Bacterial Chromatin. J Mol Biol, 431, 4040–4066.

15. Boudreau, B.A., Hron, D.R., Qin, L., van der Valk, R.A., Kotlajich, M.V., Dame, R.T. and Landick, R. (2018) StpA and Hha stimulate pausing by RNA polymerase by promoting DNA-DNA bridging of H-NS filaments. Nucleic Acids Res, 46, 5525–5546.

16. Bossi, L., Ratel, M., Laurent, C., Kerboriou, P., Camilli, A., Eveno, E., Boudvillain, M. and Figueroa-Bossi, N. (2019) NusG prevents transcriptional invasion of H-NS-silenced genes. PLoS Genet, 15, e1008425.

17. Zuber, P.K., Said, N., Hilal, T., Wang, B., Loll, B., Gonzalez-Higueras, J., Ramirez-Sarmiento, C.A., Belogurov, G.A., Artsimovitch, I., Wahl, M.C. et al. (2024) Concerted transformation of a hyper-paused transcription complex and its reinforcing protein. Nat Commun, 15, 3040.

18. Artsimovitch, I. and Landick, R. (2002) The transcriptional regulator RfaH stimulates RNA chain synthesis after recruitment to elongation complexes by the exposed nontemplate DNA strand. Cell, 109, 193–203.

19. Sevostyanova, A., Belogurov, G.A., Mooney, R.A., Landick, R. and Artsimovitch, I. (2011) The beta subunit gate loop is required for RNA polymerase modification by RfaH and NusG. Mol Cell, 43, 253–262.

20. Belogurov, G.A., Mooney, R.A., Svetlov, V., Landick, R. and Artsimovitch, I. (2009) Functional specialization of transcription elongation factors. EMBO J, 28, 112–122.

21. Burmann, B.M., Knauer, S.H., Sevostyanova, A., Schweimer, K., Mooney, R.A., Landick, R., Artsimovitch, I. and Rosch, P. (2012) An alpha helix to beta barrel domain switch transforms the transcription factor RfaH into a translation factor. Cell, 150, 291–303.

22. Hustmyer, C.M., Wolfe, M.B., Welch, R.A. and Landick, R. (2022) RfaH Counter-Silences Inhibition of Transcript Elongation by H-NS-StpA Nucleoprotein Filaments in Pathogenic Escherichia coli. mBio, 13, e0266222.

23. Wang, B., Mittermeier, M. and Artsimovitch, I. (2022) RfaH May Oppose Silencing by H-NS and YmoA Proteins during Transcription Elongation. J Bacteriol, 204, e0059921.

24. Gao, Y., Xie, N., Ma, T., Tan, C.E., Wang, Z., Zhang, R., Ma, S., Deng, Z., Wang, Y. and Shen, J. (2025) VirBR counter-silences HppX3 to promote conjugation of blaNDM-IncX3 plasmids. Nucleic Acids Res, 53.

25. Yang, J., Lu, Y., Yu, J., Cai, X., Wang, C., Lv, L., Moran, R.A., Zhao, X., Hu, Z., Deng, M. et al. (2025) Comprehensive analysis of Enterobacteriaceae IncX plasmids reveals robust conjugation regulators PrfaH, H-NS, and conjugation-fitness tradeoff. Commun Biol, 8, 363.

26. Wang, A., Cordova, M. and Navarre, W.W. (2025) Evolutionary and functional divergence of Sfx, a plasmid-encoded H-NS homolog, underlies the regulation of IncX plasmid conjugation. mBio, 16, e0208924.

27. Shi, X. and Bennett, G.N. (1994) Plasmids bearing hfq and the hns-like gene stpA complement hns mutants in modulating arginine decarboxylase gene expression in Escherichia coli. J Bacteriol, 176, 6769–6775.

28. Doyle, M., Fookes, M., Ivens, A., Mangan, M.W., Wain, J. and Dorman, C.J. (2007) An H-NS-like stealth protein aids horizontal DNA transmission in bacteria. Science, 315, 251–252.

29. Dillon, S.C., Cameron, A.D., Hokamp, K., Lucchini, S., Hinton, J.C. and Dorman, C.J. (2010) Genome-wide analysis of the H-NS and Sfh regulatory networks in Salmonella Typhimurium identifies a plasmid-encoded transcription silencing mechanism. Mol Microbiol, 76, 1250–1265.

30. Molano, L.-A.G., Hirsch, P., Hannig, M., Müller, R. and Keller, A. (2024) The PLSDB 2025 update: enhanced annotations and improved functionality for comprehensive plasmid research. Nucleic Acids Research, 53, D189–D196.

31. Camacho, C., Coulouris, G., Avagyan, V., Ma, N., Papadopoulos, J., Bealer, K. and Madden, T.L. (2009) BLAST+: architecture and applications. BMC Bioinformatics, 10, 421.

32. Emenecker, R.J., Griffith, D. and Holehouse, A.S. (2021) Metapredict: a fast, accurate, and easy-to-use predictor of consensus disorder and structure. Biophys J, 120, 4312–4319.

33. Livak, K.J. and Schmittgen, T.D. (2001) Analysis of relative gene expression data using real-time quantitative PCR and the 2(-Delta Delta C(T)) Method. Methods, 25, 402–408.

34. Adams, P.P., Baniulyte, G., Esnault, C., Chegireddy, K., Singh, N., Monge, M., Dale, R.K., Storz, G. and Wade, J.T. (2021) Regulatory roles of Escherichia coli 5’ UTR and ORF-internal RNAs detected by 3’ end mapping. eLife, 10, e62438.

35. Freddolino, L., Amemiya, H.M., Goss, T.J. and Tavazoie, S. (2021) Dynamic landscape of protein occupancy across the Escherichia coli chromosome. PLOS Biology, 19, e3001306.

36. Felix Krueger, F.J., Phil Ewels, Ebrahim Afyounian, & Benjamin Schuster-Boeckler. (2021) FelixKrueger/TrimGalore: v0.6.7 - DOI via Zenodo (0.6.7). Zenodo 10.5281/zenodo.5127899.

37. Langmead, B. and Salzberg, S.L. (2012) Fast gapped-read alignment with Bowtie 2. Nature Methods, 9, 357–359.

38. Danecek, P., Bonfield, J.K., Liddle, J., Marshall, J., Ohan, V., Pollard, M.O., Whitwham, A., Keane, T., McCarthy, S.A., Davies, R.M. et al. (2021) Twelve years of SAMtools and BCFtools. Gigascience, 10.

39. Liao, Y., Smyth, G.K. and Shi, W. (2014) featureCounts: an efficient general purpose program for assigning sequence reads to genomic features. Bioinformatics, 30, 923–930.

40. Love, M.I., Huber, W. and Anders, S. (2014) Moderated estimation of fold change and dispersion for RNA-seq data with DESeq2. Genome Biol, 15, 550.

41. Ramírez, F., Ryan, D.P., Grüning, B., Bhardwaj, V., Kilpert, F., Richter, A.S., Heyne, S., Dündar, F. and Manke, T. (2016) deepTools2: a next generation web server for deep-sequencing data analysis. Nucleic Acids Research, 44, W160–W165.

42. Zhang, Y., Liu, T., Meyer, C.A., Eeckhoute, J., Johnson, D.S., Bernstein, B.E., Nusbaum, C., Myers, R.M., Brown, M., Li, W. et al. (2008) Model-based analysis of ChIP-Seq (MACS). Genome Biol, 9, R137.

43. Heinz, S., Benner, C., Spann, N., Bertolino, E., Lin, Y.C., Laslo, P., Cheng, J.X., Murre, C., Singh, H. and Glass, C.K. (2010) Simple combinations of lineage-determining transcription factors prime cis-regulatory elements required for macrophage and B cell identities. Mol Cell, 38, 576–589.

44. Thorvaldsdóttir, H., Robinson, J.T. and Mesirov, J.P. (2012) Integrative Genomics Viewer (IGV): high-performance genomics data visualization and exploration. Briefings in Bioinformatics, 14, 178–192.

45. Kent, W.J., Zweig, A.S., Barber, G., Hinrichs, A.S. and Karolchik, D. (2010) BigWig and BigBed: enabling browsing of large distributed datasets. Bioinformatics, 26, 2204–2207.

46. Solovyev, S., Salamov, A. and Li, R. (2011). Nova Science Publishers. In.

47. Reese, M.G. (2001) Application of a time-delay neural network to promoter annotation in the Drosophila melanogaster genome. Comput Chem, 26, 51–56.

48. Stothard, P. (2000) The sequence manipulation suite: JavaScript programs for analyzing and formatting protein and DNA sequences. Biotechniques, 28, 1102, 1104.

49. Guillerez, J., Lopez, P.J., Proux, F., Launay, H. and Dreyfus, M. (2005) A mutation in T7 RNA polymerase that facilitates promoter clearance. Proc Natl Acad Sci U S A, 102, 5958–5963.

50. Schindelin, J., Arganda-Carreras, I., Frise, E., Kaynig, V., Longair, M., Pietzsch, T., Preibisch, S., Rueden, C., Saalfeld, S., Schmid, B. et al. (2012) Fiji: an open-source platform for biological-image analysis. Nature Methods, 9, 676–682.

51. Lee, C., Kim, J., Shin, S.G. and Hwang, S. (2006) Absolute and relative QPCR quantification of plasmid copy number in Escherichia coli. J Biotechnol, 123, 273–280.

52. Núñez, B., Avila, P. and de la Cruz, F. (1997) Genes involved in conjugative DNA processing of plasmid R6K. Mol Microbiol, 24, 1157–1168.

53. Borowiec, J.A. and Gralla, J.D. (1986) High-resolution analysis of lac transcription complexes inside cells. Biochemistry, 25, 5051–5057.

54. Yang, J., Lu, Y., Yu, J., Cai, X., Wang, C., Lv, L., Moran, R.A., Zhao, X., Hu, Z., Deng, M. et al. (2025) Comprehensive analysis of Enterobacteriaceae IncX plasmids reveals robust conjugation regulators PrfaH, H-NS, and conjugation-fitness tradeoff. Communications Biology, 8, 363.

55. Gao, Y., Xie, N., Ma, T., Tan, Chun E., Wang, Z., Zhang, R., Ma, S., Deng, Z., Wang, Y. and Shen, J. (2025) VirBR counter-silences HppX3 to promote conjugation of blaNDM-IncX3 plasmids. Nucleic Acids Research, 53.

56. Saxena, S. and Gowrishankar, J. (2011) Modulation of Rho-dependent transcription termination in Escherichia coli by the H-NS family of proteins. J Bacteriol, 193, 3832–3841.

57. Singh, S.S., Singh, N., Bonocora, R.P., Fitzgerald, D.M., Wade, J.T. and Grainger, D.C. (2014) Widespread suppression of intragenic transcription initiation by H-NS. Genes Dev, 28, 214–219.

58. Kotlajich, M.V., Hron, D.R., Boudreau, B.A., Sun, Z., Lyubchenko, Y.L. and Landick, R. (2015) Bridged filaments of histone-like nucleoid structuring protein pause RNA polymerase and aid termination in bacteria. eLife, 4, e04970.

59. Matsumoto, Y., Shigesada, K., Hirano, M. and Imai, M. (1986) Autogenous regulation of the gene for transcription termination factor rho in Escherichia coli: localization and function of its attenuators. J Bacteriol, 166, 945–958.

60. Chant, E.L. and Summers, D.K. (2007) Indole signalling contributes to the stable maintenance of Escherichia coli multicopy plasmids. Mol Microbiol, 63, 35–43.

61. van der Stel, A.X., Gordon, E.R., Sengupta, A., Martínez, A.K., Klepacki, D., Perry, T.N., Herrero Del Valle, A., Vázquez-Laslop, N., Sachs, M.S., Cruz-Vera, L.R., et al. (2021) Structural basis for the tryptophan sensitivity of TnaC-mediated ribosome stalling. Nat Commun, 12, 5340.

62. Stewart, V. and Yanofsky, C. (1985) Evidence for transcription antitermination control of tryptophanase operon expression in Escherichia coli K-12. J Bacteriol, 164, 731–740.

63. Gasperotti, A., Göing, S., Fajardo-Ruiz, E., Forné, I. and Jung, K. (2020) Function and Regulation of the Pyruvate Transporter CstA in Escherichia coli. Int J Mol Sci, 21.

64. Qiu, J., Gasperotti, A., Sisattana, N., Zacharias, M. and Jung, K. (2023) The LytS-type histidine kinase BtsS is a 7-transmembrane receptor that binds pyruvate. mBio, 14, e0108923.

65. Behr, S., Kristoficova, I., Witting, M., Breland, E.J., Eberly, A.R., Sachs, C., Schmitt-Kopplin, P., Hadjifrangiskou, M. and Jung, K. (2017) Identification of a High-Affinity Pyruvate Receptor in Escherichia coli. Sci Rep, 7, 1388.

66. Dorman, C.J. (2004) H-NS: a universal regulator for a dynamic genome. Nature Reviews Microbiology, 2, 391–400.

67. Wang, A., Cordova, M. and Navarre, W.W. (2025) Evolutionary and functional divergence of Sfx, a plasmid-encoded H-NS homolog, underlies the regulation of IncX plasmid conjugation. mBio, 16, e02089–02024.

68. Uyar, E., Kurokawa, K., Yoshimura, M., Ishikawa, S., Ogasawara, N. and Oshima, T. (2009) Differential binding profiles of StpA in wild-type and h-ns mutant cells: a comparative analysis of cooperative partners by chromatin immunoprecipitation-microarray analysis. J Bacteriol, 191, 2388–2391.

69. Kahramanoglou, C., Seshasayee, A.S., Prieto, A.I., Ibberson, D., Schmidt, S., Zimmermann, J., Benes, V., Fraser, G.M. and Luscombe, N.M. (2011) Direct and indirect effects of H-NS and Fis on global gene expression control in Escherichia coli. Nucleic Acids Res, 39, 2073–2091.

70. Kontomichalou, P., Mitani, M. and Clowes, R.C. (1970) Circular R-factor molecules controlling penicillinase synthesis, replicating in Escherichia coli under either relaxed or stringent control. J Bacteriol, 104, 34–44.

71. Free, A. and Dorman, C.J. (1995) Coupling of Escherichia coli hns mRNA levels to DNA synthesis by autoregulation: implications for growth phase control. Mol Microbiol, 18, 101–113.

72. Navarre, W.W., Porwollik, S., Wang, Y., McClelland, M., Rosen, H., Libby, S.J. and Fang, F.C. (2006) Selective silencing of foreign DNA with low GC content by the H-NS protein in Salmonella. Science, 313, 236–238.

73. Gordon, B.R.G., Li, Y., Cote, A., Weirauch, M.T., Ding, P., Hughes, T.R., Navarre, W.W., Xia, B. and Liu, J. (2011) Structural basis for recognition of AT-rich DNA by unrelated xenogeneic silencing proteins. Proceedings of the National Academy of Sciences, 108, 10690–10695.

74. Fang, F.C. and Rimsky, S. (2008) New insights into transcriptional regulation by H-NS. Current Opinion in Microbiology, 11, 113–120.

75. Lang, B., Blot, N., Bouffartigues, E., Buckle, M., Geertz, M., Gualerzi, C.O., Mavathur, R., Muskhelishvili, G., Pon, C.L., Rimsky, S. et al. (2007) High-affinity DNA binding sites for H-NS provide a molecular basis for selective silencing within proteobacterial genomes. Nucleic Acids Res, 35, 6330–6337.

76. Zhang, Y., Liu, T., Meyer, C.A., Eeckhoute, J., Johnson, D.S., Bernstein, B.E., Nusbaum, C., Myers, R.M., Brown, M., Li, W. et al. (2008) Model-based Analysis of ChIP-Seq (MACS). Genome Biology, 9, R137.

77. Figueroa-Bossi, N., Fernández-Fernández, R., Kerboriou, P., Bouloc, P., Casadesús, J., Sánchez-Romero, M.A. and Bossi, L. (2024) Transcription-driven DNA supercoiling counteracts H-NS-mediated gene silencing in bacterial chromatin. Nature Communications, 15, 2787.

78. Arold, S.T., Leonard, P.G., Parkinson, G.N. and Ladbury, J.E. (2010) H-NS forms a superhelical protein scaffold for DNA condensation. Proc Natl Acad Sci U S A, 107, 15728–15732.

79. Picker, M.A., Karney, M.M.A., Gerson, T.M., Karabachev, A.D., Duhart, J.C., McKenna, J.A. and Wing, H.J. (2023) Localized modulation of DNA supercoiling, triggered by the Shigella anti-silencer VirB, is sufficient to relieve H-NS-mediated silencing. Nucleic Acids Res, 51, 3679–3695.

80. Spassky, A., Rimsky, S., Garreau, H. and Buc, H. (1984) H1a, an E. coli DNA-binding protein which accumulates in stationary phase, strongly compacts DNA in vitro. Nucleic Acids Res, 12, 5321–5340.

81. Tupper, A.E., Owen-Hughes, T.A., Ussery, D.W., Santos, D.S., Ferguson, D.J., Sidebotham, J.M., Hinton, J.C. and Higgins, C.F. (1994) The chromatin-associated protein H-NS alters DNA topology in vitro. EMBO J, 13, 258–268.

82. Lal, A., Dhar, A., Trostel, A., Kouzine, F., Seshasayee, A.S. and Adhya, S. (2016) Genome scale patterns of supercoiling in a bacterial chromosome. Nat Commun, 7, 11055.

83. Guo, M.S., Kawamura, R., Littlehale, M.L., Marko, J.F. and Laub, M.T. (2021) High-resolution, genome-wide mapping of positive supercoiling in chromosomes. Elife, 10.

84. Sinden, R.R., Carlson, J.O. and Pettijohn, D.E. (1980) Torsional tension in the DNA double helix measured with trimethylpsoralen in living E. coli cells: analogous measurements in insect and human cells. Cell, 21, 773–783.

85. Gellert, M., O’Dea, M.H., Itoh, T. and Tomizawa, J. (1976) Novobiocin and coumermycin inhibit DNA supercoiling catalyzed by DNA gyrase. Proc Natl Acad Sci U S A, 73, 4474–4478.

86. Dole, S., Nagarajavel, V. and Schnetz, K. (2004) The histone-like nucleoid structuring protein H-NS represses the Escherichia coli bgl operon downstream of the promoter. Mol Microbiol, 52, 589–600.

87. Brescia, C.C., Kaw, M.K. and Sledjeski, D.D. (2004) The DNA binding protein H-NS binds to and alters the stability of RNA in vitro and in vivo. J Mol Biol, 339, 505–514.

88. Mayer, O., Rajkowitsch, L., Lorenz, C., Konrat, R. and Schroeder, R. (2007) RNA chaperone activity and RNA-binding properties of the E. coli protein StpA. Nucleic Acids Res, 35, 1257–1269.

89. Park, H.S., Ostberg, Y., Johansson, J., Wagner, E.G. and Uhlin, B.E. (2010) Novel role for a bacterial nucleoid protein in translation of mRNAs with suboptimal ribosome-binding sites. Genes Dev, 24, 1345–1350.

90. Johansson, J., Dagberg, B., Richet, E. and Uhlin, B.E. (1998) H-NS and StpA proteins stimulate expression of the maltose regulon in Escherichia coli. J Bacteriol, 180, 6117–6125.

91. Gavrilov, A.A., Shamovsky, I., Zhegalova, I., Proshkin, S., Shamovsky, Y., Evko, G., Epshtein, V., Rasouly, A., Blavatnik, A., Lahiri, S. et al. (2025) Elementary 3D organization of active and silenced E. coli genome. Nature, 645, 1060–1070.

92. Amemiya, H.M., Schroeder, J. and Freddolino, P.L. (2021) Nucleoid-associated proteins shape chromatin structure and transcriptional regulation across the bacterial kingdom. Transcription, 12, 182–218.

93. Kaval, K.G., Chimalapati, S., Siegel, S.D., Garcia, N., Jaishankar, J., Dalia, A.B. and Orth, K. (2023) Membrane-localized expression, production and assembly of Vibrio parahaemolyticus T3SS2 provides evidence for transertion. Nature Communications, 14, 1178.

94. Libby, E.A., Roggiani, M. and Goulian, M. (2012) Membrane protein expression triggers chromosomal locus repositioning in bacteria. Proc Natl Acad Sci U S A, 109, 7445–7450.

95. Lodge, J.K., Kazic, T. and Berg, D.E. (1989) Formation of supercoiling domains in plasmid pBR322. J Bacteriol, 171, 2181–2187.

96. Li, Z., Li, Z., Peng, Y., Zhang, M., Wen, Y., Lu, X. and Kan, B. (2025) Genomic diversity of mcr-carrying plasmids and the role of type IV secretion systems in IncI2 plasmids conjugation. Commun Biol, 8, 342.

97. Mace, K., Vadakkepat, A.K., Redzej, A., Lukoyanova, N., Oomen, C., Braun, N., Ukleja, M., Lu, F., Costa, T.R.D., Orlova, E.V. et al. (2022) Cryo-EM structure of a type IV secretion system. Nature, 607, 191–196.

98. Arutyunov, D. and Frost, L.S. (2013) F conjugation: back to the beginning. Plasmid, 70, 18–32.

99. Garcillan-Barcia, M.P. and de la Cruz, F. (2008) Why is entry exclusion an essential feature of conjugative plasmids? Plasmid, 60, 1–18.

100. Audette, G.F., Manchak, J., Beatty, P., Klimke, W.A. and Frost, L.S. (2007) Entry exclusion in F-like plasmids requires intact TraG in the donor that recognizes its cognate TraS in the recipient. Microbiology (Reading), 153, 442–451.

101. Humbert, M., Huguet, K.T., Coulombe, F. and Burrus, V. (2019) Entry Exclusion of Conjugative Plasmids of the IncA, IncC, and Related Untyped Incompatibility Groups. J Bacteriol, 201.

102. Vlijm, R., J, V.D.T. and Dekker, C. (2015) Counterintuitive DNA Sequence Dependence in Supercoiling-Induced DNA Melting. PLoS One, 10, e0141576.

103. Oberstrass, F.C., Fernandes, L.E., Lebel, P. and Bryant, Z. (2013) Torque spectroscopy of DNA: base-pair stability, boundary effects, backbending, and breathing dynamics. Phys Rev Lett, 110, 178103.

104. Martin, E.W. and Holehouse, A.S. (2020) Intrinsically disordered protein regions and phase separation: sequence determinants of assembly or lack thereof. Emerg Top Life Sci, 4, 307–329.

105. Hirose, T., Ninomiya, K., Nakagawa, S. and Yamazaki, T. (2023) A guide to membraneless organelles and their various roles in gene regulation. Nature Reviews Molecular Cell Biology, 24, 288–304.

106. Müller, C.M., Dobrindt, U., Nagy, G., Emödy, L., Uhlin, B.E. and Hacker, J. (2006) Role of histone-like proteins H-NS and StpA in expression of virulence determinants of uropathogenic Escherichia coli. J Bacteriol, 188, 5428–5438.

107. Gehrke, E.J., Zhang, X., Pimentel-Elardo, S.M., Johnson, A.R., Rees, C.A., Jones, S.E., Hindra, Gehrke, S.S., Turvey, S., Boursalie, S., et al. (2019) Silencing cryptic specialized metabolism in Streptomyces by the nucleoid-associated protein Lsr2. Elife, 8.

108. Ramirez, V., Tiller, L., Szafran, M.J., Zhang, X., Jakimowicz, D., Elliot, M. and Freddolino, L. (2025) LsrL modulates Lsr2-induced chromatin structure to tune biosynthetic gene cluster regulation in Streptomyces venezuelae. bioRxiv.

109. Castang, S., McManus, H.R., Turner, K.H. and Dove, S.L. (2008) H-NS family members function coordinately in an opportunistic pathogen. Proceedings of the National Academy of Sciences, 105, 18947–18952.

110. Li, C., Wally, H., Miller, S.J. and Lu, C.-D. (2009) The Multifaceted Proteins MvaT and MvaU, Members of the H-NS Family, Control Arginine Metabolism, Pyocyanin Synthesis, and Prophage Activation in *Pseudomonas aeruginosa* PAO1. Journal of Bacteriology, 191, 6211–6218.

111. Fitzgerald, S., Kary, S.C., Alshabib, E.Y., MacKenzie, K.D., Stoebel, Daniel M., Chao, T.-C. and Cameron, A.D.S. (2020) Redefining the H-NS protein family: a diversity of specialized core and accessory forms exhibit hierarchical transcriptional network integration. Nucleic Acids Research, 48, 10184–10198.

112. Figueroa-Bossi, N., Sanchez-Romero, M.A., Kerboriou, P., Naquin, D., Mendes, C., Bouloc, P., Casadesus, J. and Bossi, L. (2022) Pervasive transcription enhances the accessibility of H-NS-silenced promoters and generates bistability in Salmonella virulence gene expression. Proc Natl Acad Sci U S A, 119, e2203011119.

113. Ishihama, A. and Shimada, T. (2021) Hierarchy of transcription factor network in Escherichia coli K-12: H-NS-mediated silencing and Anti-silencing by global regulators. FEMS Microbiol Rev, 45.

114. Doetsch, M., Gstrein, T., Schroeder, R. and Furtig, B. (2010) Mechanisms of StpA-mediated RNA remodeling. RNA Biol, 7, 735–743.

115. Waldsich, C., Grossberger, R. and Schroeder, R. (2002) RNA chaperone StpA loosens interactions of the tertiary structure in the td group I intron in vivo. Genes Dev, 16, 2300–2312.

116. Lim, C.J., Whang, Y.R., Kenney, L.J. and Yan, J. (2011) Gene silencing H-NS paralogue StpA forms a rigid protein filament along DNA that blocks DNA accessibility. Nucleic Acids Research, 40, 3316–3328.

117. Horiuchi, T. and Hidaka, M. (1988) Core sequence of two separable terminus sites of the R6K plasmid that exhibit polar inhibition of replication is a 20 bp inverted repeat. Cell, 54, 515–523.

118. Lovett, M.A., Sparks, R.B. and Helinski, D.R. (1975) Bidirectional replication of plasmid R6K DNA in Escherichia coli; correspondence between origin of replication and position of single-strand break in relaxed complex. Proc Natl Acad Sci U S A, 72, 2905–2909.

119. Lim, C.J., Kenney, L.J. and Yan, J. (2014) Single-molecule studies on the mechanical interplay between DNA supercoiling and H-NS DNA architectural properties. Nucleic Acids Res, 42, 8369–8378.

120. Kim, S.H., Ganji, M., Kim, E., van der Torre, J., Abbondanzieri, E. and Dekker, C. (2018) DNA sequence encodes the position of DNA supercoils. Elife, 7.

121. Lukose, B., Goyal, S. and Naganathan, A.N. (2025) Oligomerization-mediated phase separation in the nucleoid-associated sensory protein H-NS is controlled by ambient cues. Protein Sci, 34, e5250.

122. Noom, M.C., Navarre, W.W., Oshima, T., Wuite, G.J. and Dame, R.T. (2007) H-NS promotes looped domain formation in the bacterial chromosome. Curr Biol, 17, R913–914.

123. Shen, B.A., Hustmyer, C.M., Roston, D., Wolfe, M.B. and Landick, R. (2022) Bacterial H-NS contacts DNA at the same irregularly spaced sites in both bridged and hemi-sequestered linear filaments. iScience, 25.

124. Schmidt, A., Kochanowski, K., Vedelaar, S., Ahrne, E., Volkmer, B., Callipo, L., Knoops, K., Bauer, M., Aebersold, R. and Heinemann, M. (2016) The quantitative and condition-dependent Escherichia coli proteome. Nat Biotechnol, 34, 104–110.

125. Ali, S.S., Whitney, J.C., Stevenson, J., Robinson, H., Howell, P.L. and Navarre, W.W. (2013) Structural insights into the regulation of foreign genes in Salmonella by the Hha/H-NS complex. J Biol Chem, 288, 13356–13369.

126. Abramson, J., Adler, J., Dunger, J., Evans, R., Green, T., Pritzel, A., Ronneberger, O., Willmore, L., Ballard, A.J., Bambrick, J. et al. (2024) Accurate structure prediction of biomolecular interactions with AlphaFold 3. Nature, 630, 493–500.

127. Ali Azam, T., Iwata, A., Nishimura, A., Ueda, S. and Ishihama, A. (1999) Growth phase-dependent variation in protein composition of the Escherichia coli nucleoid. J Bacteriol, 181, 6361–6370.

128. McCarthy, L.A., Way, L.E., Dai, X., Ren, Z., Fuller, D.E.H., Dhiman, I., Larkin, L., Sieben, J.J.D., Westerlaken, I., Abbondanzieri, E.A. et al. (2026) Dps binds and protects DNA in starved Escherichia coli with minimal effect on chromosome accessibility, dynamics, and organization. Nucleic Acids Res, 54.

129. Amemiya, H.M., Goss, T.J., Nye, T.M., Hurto, R.L., Simmons, L.A. and Freddolino, P.L. (2022) Distinct heterochromatin-like domains promote transcriptional memory and silence parasitic genetic elements in bacteria. EMBO J, 41, e108708.

130. Kuznetsova, M.V., Gizatullina, J.S., Koraimann, G. and Erjavec, M.S. (2022) The prospect of creating farm animal probiotics based on genetically modified Escherichia coli strains. AIP Conference Proceedings, 2390.

131. Decker, K.T., Gao, Y., Rychel, K., Al Bulushi, T., Chauhan, Siddharth M., Kim, D., Cho, B.-K. and Palsson, Bernhard O. (2021) proChIPdb: a chromatin immunoprecipitation database for prokaryotic organisms. Nucleic Acids Research, 50, D1077–D1084.

