## Supplementary figures and table for "Selective targeting of a histone-like silencer Sfx to the R6K conjugal transfer operon"

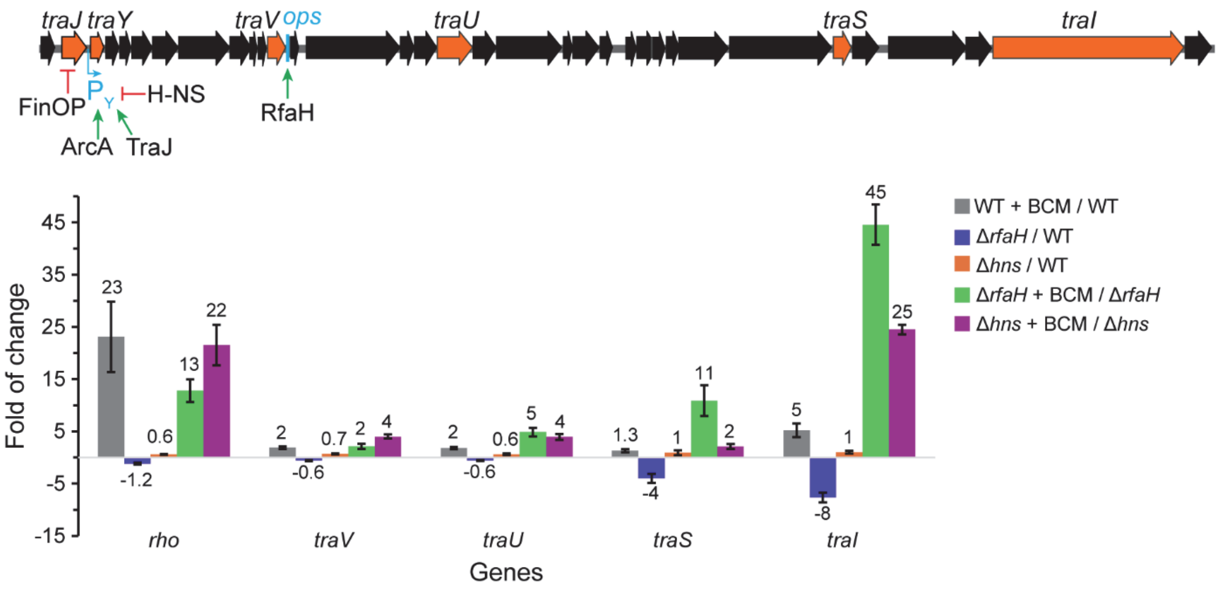


**Supplementary Figure S1**. The effects of Rho, RfaH, and H-NS on the F-factor *tra* operon were tested using RT-qPCR. WT, wild-type *E. coli* strain. BCM, bicyclomycin. Mean fold changes are indicated on top of the bars. Negative values represent reduced expression. Error bars represent the SD of three biological repeats.

F is a single-copy 100 kb plasmid that carries the 33-kb *tra* operon transcribed from the $P_Y$ promoter. $P_Y$ is activated by the F-encoded TraJ and host co-activator ArcA (1) and repressed by H-NS (2). FinOP represses the translation of TraJ, but FinOP was disrupted by an IS3 insertion in F plasmid (3). The post-initiation control of the *tra* operon, the longest operon in *E. coli*, has not been investigated, but is suggested by an early report that its expression depends on RfaH (4), a host antitermination factor required for the expression of other long horizontally acquired operons that contain an *ops* DNA element (5).

In F, the *ops* element is located downstream from the *traV* (*virB7*) gene, potentially placing the downstream genes under RfaH control. Indeed, we found that the deletion of *rfaH* reduced the expression of distal *traS* and *traI* genes by 4- and 8-fold. The observed polarity was reversed in the presence of BCM, a specific inhibitor of Rho; the *traI* expression in the *ΔrfaH* strain was increased 45-fold when Rho was inhibited by BCM. The *hns* deletion did not alleviate polarity of the derepressed F-factor when RfaH and Rho were active, but increased *traI* RNA levels ~25-fold upon the addition of BCM. These results demonstrate that the host-encoded H-NS, Rho, and RfaH modulate RNA chain elongation in the F plasmid *tra* operon, a pattern consistent with findings in other horizontally acquired *E. coli* operons (6-8). The *ops* element is present in all F-like plasmids (3), suggesting that this mode of regulation is widespread.

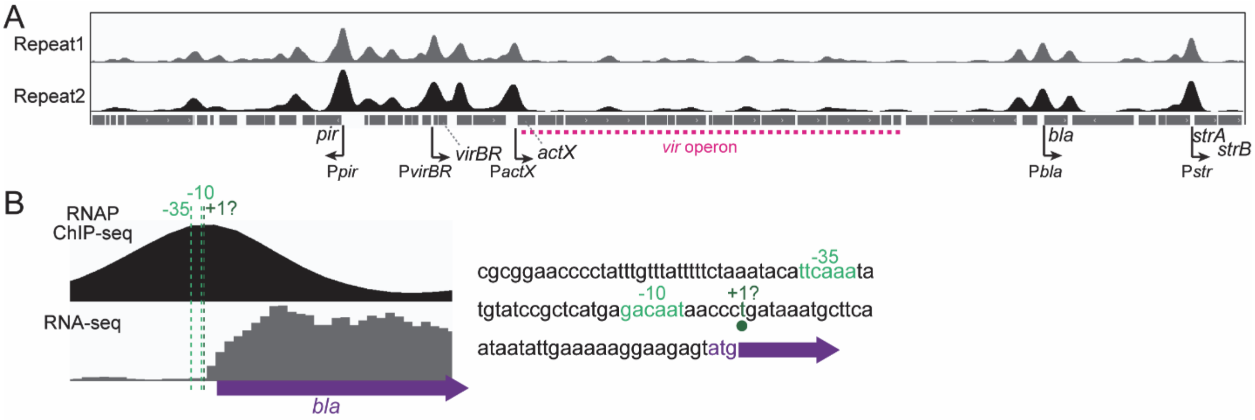


**Supplementary Figure S2**. RNAP ChIP-seq maps promoters in R6K. (**A**) The cell cultures harboring R6KΔ*sfx* were treated with rifampicin to enrich for promoter-bound RNAPs. The IGV tracks of two biological repeats, showing the fold enrichment (ranging from 0 to 10; FE = ChIP / Input), are displayed. The *pir* gene promoter has the highest RNAP peak, suggesting strong transcription. The *virBR* gene is co-transcribed with *hyp12*. The *bla* gene promoter overlaps with the known promoter region. The other two antibiotic resistance genes, *strA* and *strB*, are transcribed from the same promoter. No significant RNAP signals were detected inside the *vir* operon. (**B**) The published (9) *bla* promoter -10 and -35 elements are located around the RNAP ChIP-seq peak. The transcription start site is predicted by BDGP (10). RNA-seq coverage smoothed in 20-bp windows agrees with the locations of the promoter.

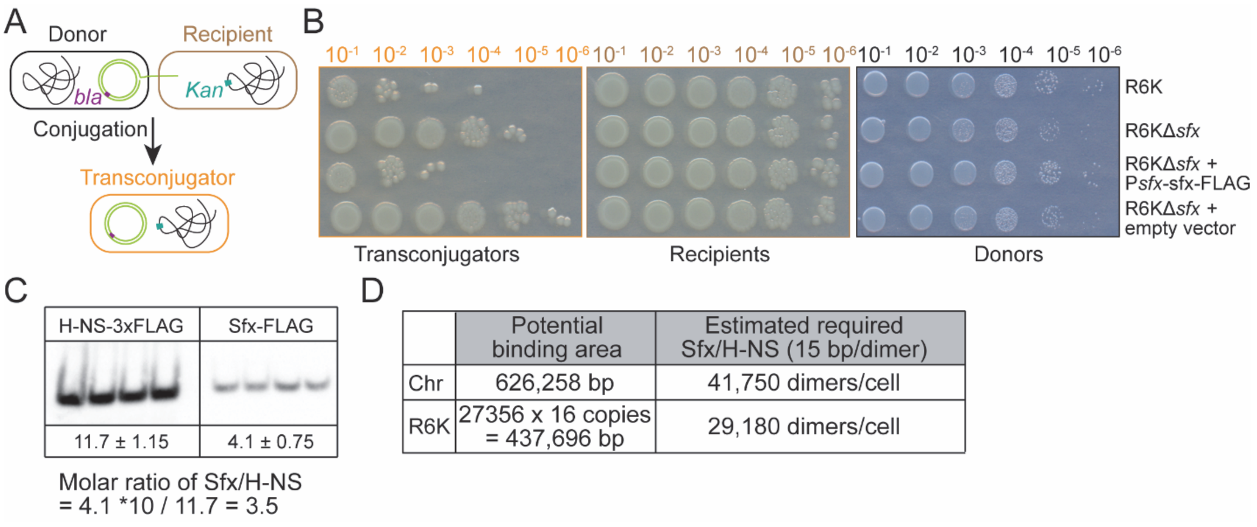


**Supplementary Figure S3**. Testing the activity and abundance of FLAG-tagged Sfx. (**A**) Schematic of conjugation assay. (**B**) Sfx-FLAG can inhibit conjugation *in trans*. (**C**) Determine H-NS/Sfx molar ratio by Western Blot. Four biological repeats are shown. The data (mean ± SD) represent the relative abundance of the target protein. The western blot signal is normalized by the total input protein for each lane, quantified by Coomassie Blue staining of the SDS-PAGE gel. According to the technical notes from Sigma, 3xFLAG has at least ten times the detection sensitivity of a single FLAG. Thus, we converted the relative abundance by a factor of ten to calculate the molar ratio. (**D**) Potential binding areas on the chromosome and R6K are estimated from the binding peaks of H-NS on the chromosome and binding peaks of Sfx on R6K, respectively. The required numbers of Sfx and H-NS dimers are estimated assuming that both proteins cover ~15 bp of DNA/dimer (11).

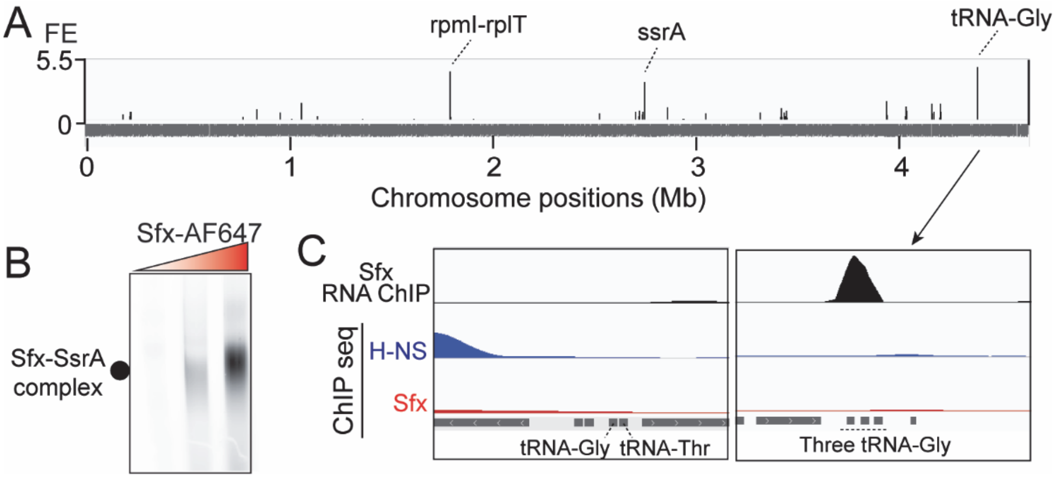


**Supplementary Figure S4**. RNA binding by Sfx. (**A**) RNA was ChIP-ed by Sfx-FLAG. Mean fold enrichment of two biological repeats is shown (FE = ChIP / Input). The top three enriched RNAs on the chromosome are indicated; there was no enriched (FE > 2 in both biological repeats) RNA on R6K. (**B**) EMSA with 100 nM *ssrA* RNA and 1, 2, and 4 μM of Alexa Fluor 647 (AF647) labeled Sfx. The Sfx-SsrA complex is visualized with AF647 fluorescence. (**C**) Track views of significantly downregulated chromosomal genes upon the *sfx* deletion. FE of H-NS and Sfx ChIP-seq tracks ranged from 0 to 10, and FE of RNA ChIP-seq data ranged from 0 to 5.5. The downregulated upon the *sfx* deletion tRNA-Gly (BW25113_RS20640) and tRNA-Thr (BW25113_RS20645) have no Sfx bound to their coding regions or the tRNAs themselves. In the RNA ChIP-seq data, we can see three enriched tRNA-Gly (BW25113_RS21635, BW25113_RS21640, BW25113_RS21645), but their RNA levels determined by RNA-seq are not changed.

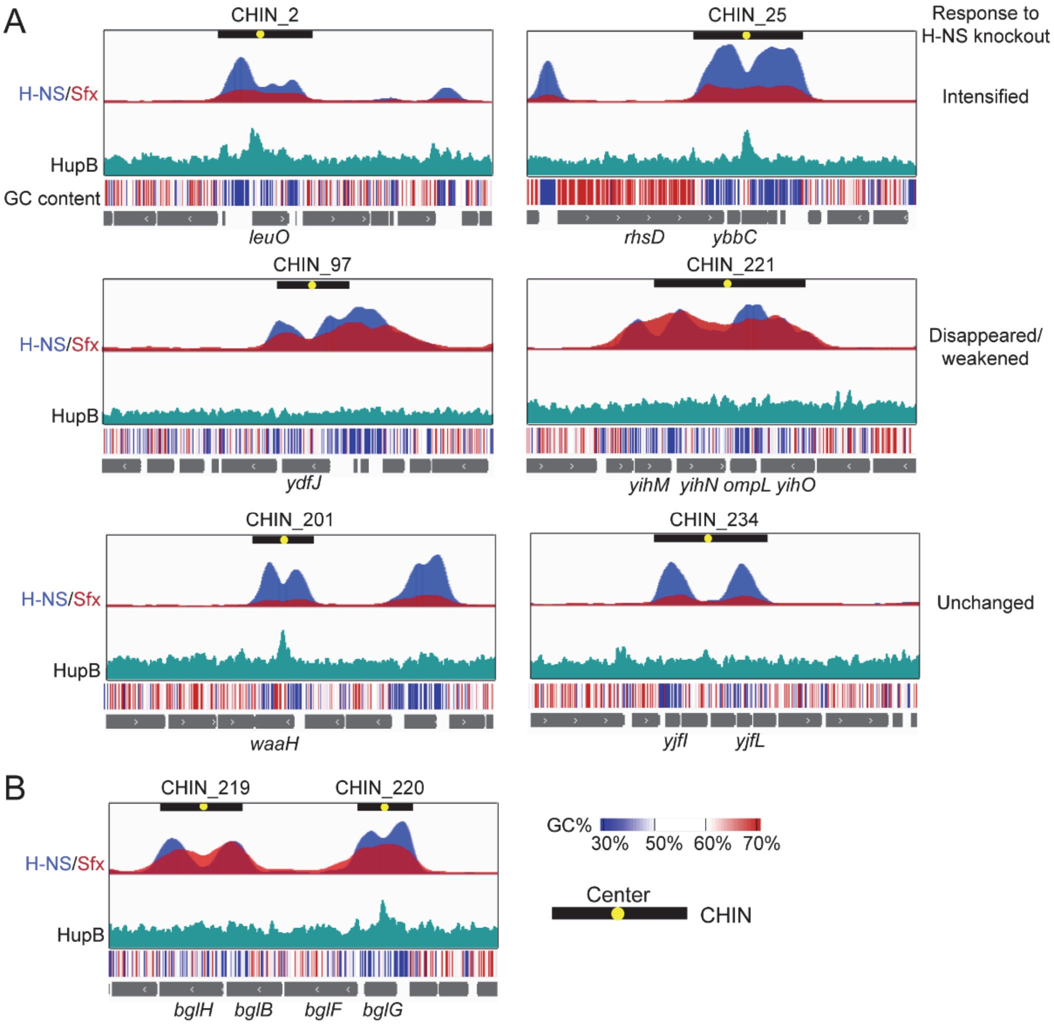


**Supplementary Figure S5**. Sfx can form continuous peaks through the CHIN loops. (**A**) ChIP-seq tracks show examples of CHINs. CHIN centers (loops) are located in the valley of H-NS binding peaks. According to the response to H-NS knockout, CHINs are separated into three groups (12), and two examples for each group are shown here. HupB binding is different in the three CHIN groups. (**B**) The *bgl* operon has two CHINs: CHIN_219 (disappeared/weakened upon H-NS knockout) doesn't have HupB peak, while the other one CHIN_220 is intensified and has HupB peak. In all observed cases, Sfx goes through the H-NS peak valley, suggesting Sfx can compete with HU for binding the loop area of CHINs. The GC content, calculated in a 25 bp sliding window, is shown as a heatmap.

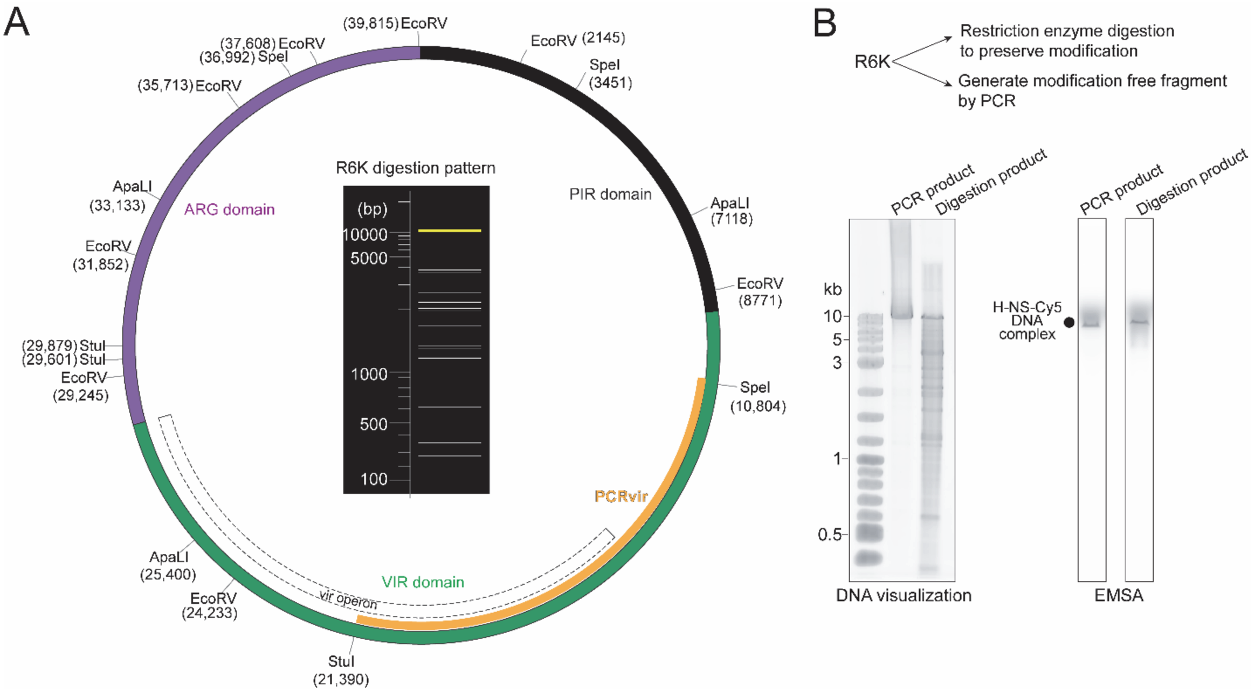


**Supplementary Figure S6**. DNA modification cannot explain the H-NS exclusion from the *vir* operon. (**A**) R6K digestion map. The R6K digestion pattern is generated by NEBcutter v3.0.23. The three R6K ARG, PIR, and VIR domains are colored in purple, black, and green, respectively. The tested DNA fragment (PCRvir) is indicated in orange. (**B**) DNA fragments containing the *vir* operon (PCRvir in panel **A** were generated by PCR (no modification) or digestion with *Apa*LI, *Eco*RV, *Spe*I, and *Stu*I restriction enzymes (to preserve potential DNA modifications) and used for EMSA with Cy5-labeled H-NS. To visualize the DNA fragments, 300 ng of DNA were stained with SYBR Gold Nucleic Acid Gel Stain (ThermoFisher). The 10.6 kb PCRvir fragment has similar intensity, quantified with ImageJ, in both PCR and digestion products (Left). One μM of H-NS-Cy5 was incubated with 4 ng/μl PCR product or digestion product for 15 min at room temperature, and the reactions were analyzed on 1% agarose gel at 100 V, in 0.5x TBE. After running the gel for 2 h at 4 °C, the gel is imaged with Amersham Typhoon 5 (Right). Although many fragments are present in the digestion product, the H-NS-Cy5 + PCRvir fragment is the single strong band. The band intensity representing DNA-bound H-NS-Cy5 was quantified, and the ratio of band intensity in the PCR product to that in the digestion product is 1.14 ± 0.16 (Mean ± SD; n = 3).

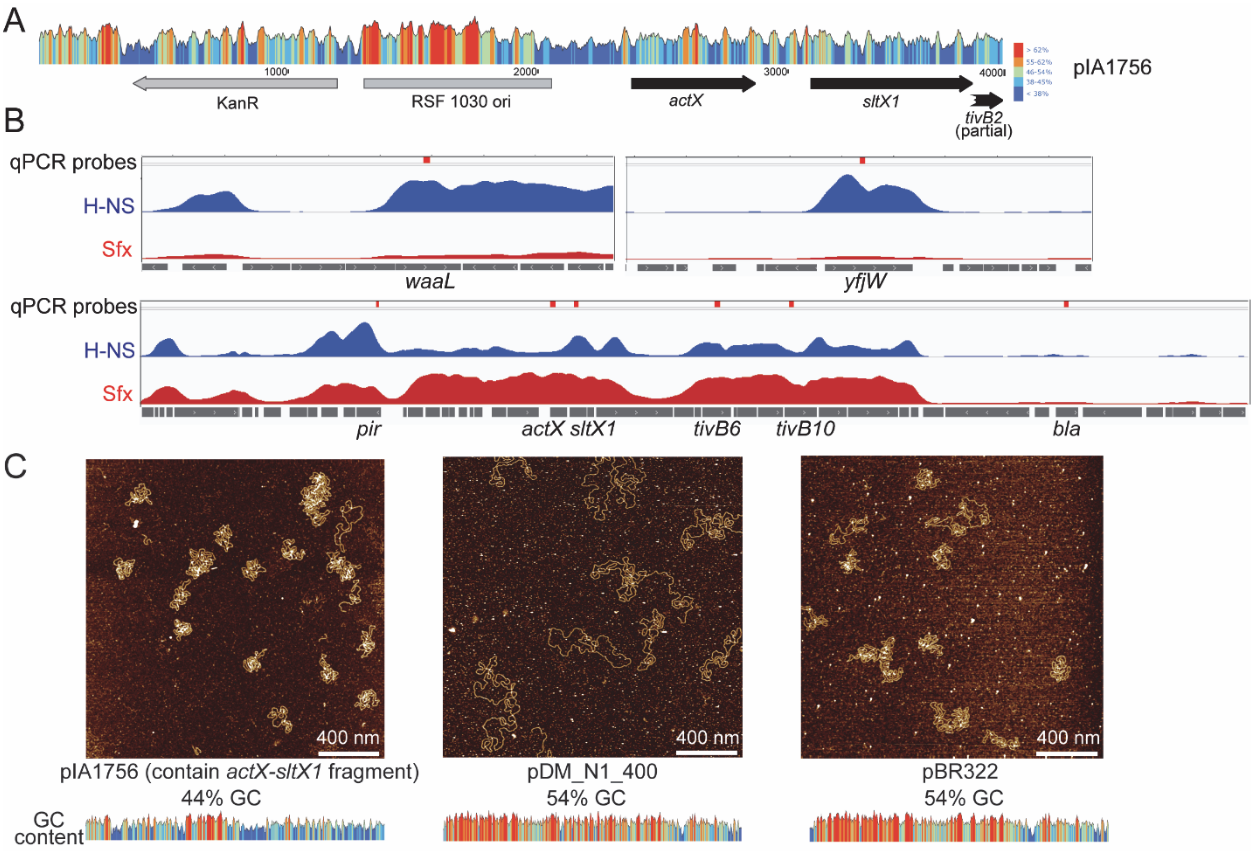


**Supplementary Figure S7**. Shuffling the *actX-sltX1* fragment. (**A**) plasmid map of pIA1756. The *actX-sltX1* transcription is under the control of Sfx. RT-qPCR shows 90 ± 9.9 (n = 3) folds of suppression in the presence of Sfx. GC content (calculated with Snapgene v8.2.2) is shown above the genes. (**B**) Positions of qPCR probes (red bars) are indicated above the ChIP-seq peaks. ChIP-seq tracks are the mean value of two biological repeats, showing the fold enrichment (ranging from 0 to 13). FE = ChIP / Input. (**C**) AFM images of pIA1756 containing the R6K *actX-sltX1* fragment, a plasmid with pBR322 backbone (pDM_N1_400), and the pBR322 plasmid. The scale bar is 400 nm. GC content distributions are shown below the images (same scale as panel **A**).

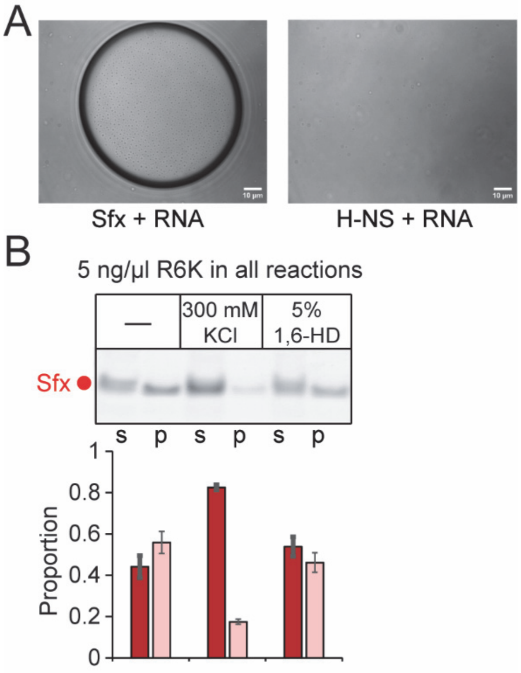


**Supplementary Figure S8**. Sfx phase separation is sensitive to salt concentration. (**A**) 5 µM Sfx forms droplets with 30 ng/µl total RNA extracted from *E. coli* in the presence of 8% PEG8000, but H-NS doesn't under the same condition. (**B**) Phase separation formation was challenged with 300 mM KCl or 5% (w/v) 1,6-Hexanediol (1,6-HD). The pelleting assay was performed as **Fig. 6B**.

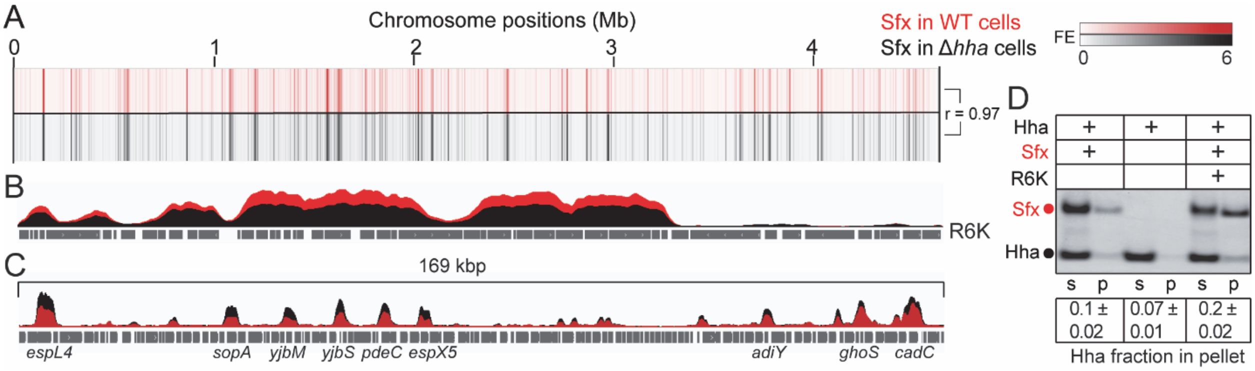


**Supplementary Figure S9**. ChIP-seq tracks of Sfx in wild-type and Δ*hha E. coli* cells. (**A**) Heatmap tracks on the chromosome. Spearman correlation (r) is calculated by binning reads at 300 bp. (**B**) The peak heights of Sfx on R6K is decreased in the absence of Hha. The peaks represent FE (fold enrichment) in the range of 0 to 11. (**C**) A 169 kbp region of the chromosome shows that Sfx peaks after the *hha* deletion become higher, possibly reflecting the decreased binding of H-NS. (**D**) A pelleting assay showing that Hha co-sediments with Sfx. In the reaction, 2 µM Sfx was mixed with 5 µM Hha and 5 ng/µl R6K, where indicated. After incubating for 10 min at room temperature, the reactions were spun at 21,000 x *g* for 25 min at 20 °C. s, supernatant. p, pellet. The Hha fraction in the pellet is presented as Mean ± SD (n = 3). Using band intensity, it's estimated that the molar ratio of Sfx/Hha in the pellet is 1, which is similar to the H-NS/Hha complex (13).

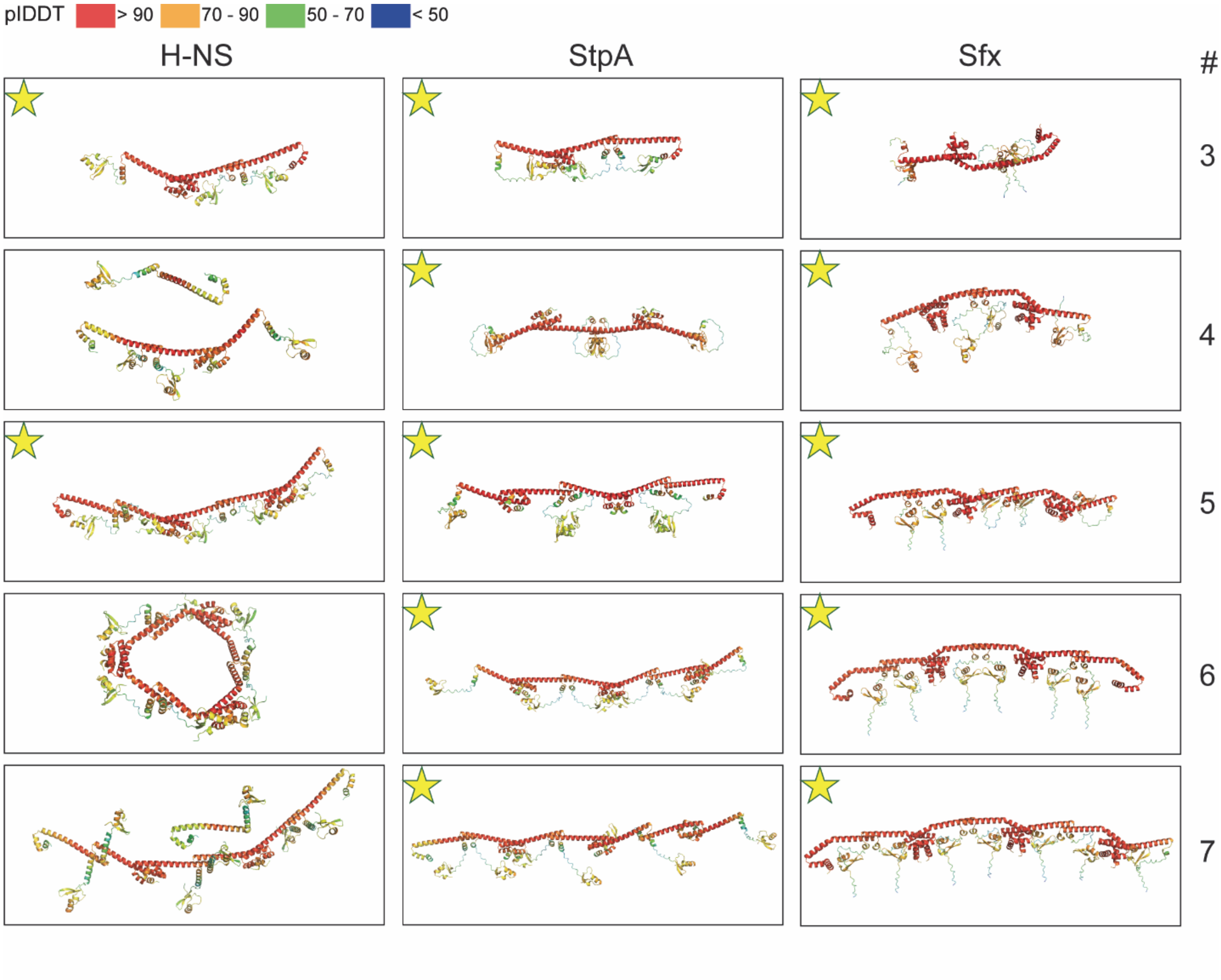


**Supplementary Figure S10**. Multimer prediction by AlphaFold3. The gold stars indicate the formation of undisrupted oligomers. The numbers on the right indicate the number of molecules used for prediction. The results show that Sfx and StpA can form oligomers in all conditions, while only two worked for H-NS.

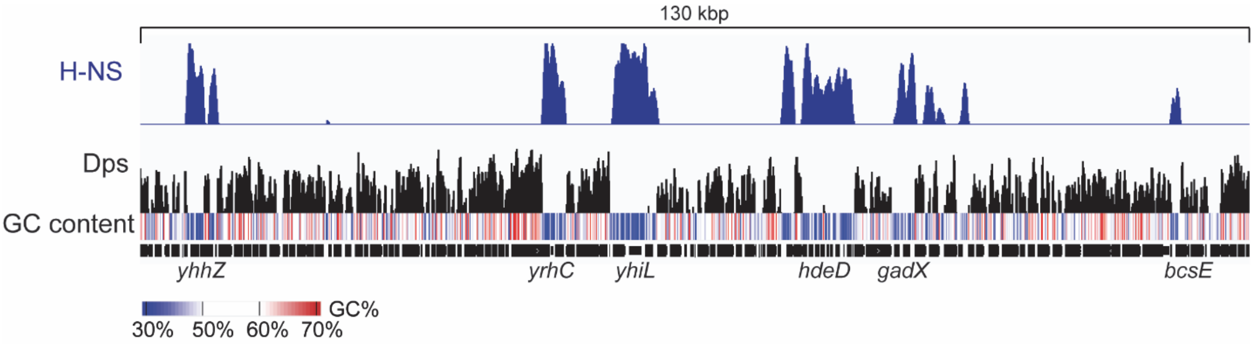


**Supplementary Figure S11**. ChIP-seq tracks of H-NS (this work) and Dps from the stationary phase (GSE293552. 24-hour samples, see reference (14)). A 130 kbp region of the chromosome demonstrates that Dps binds to high GC regions and is excluded from H-NS binding regions. The tracks represent FE (fold enrichment). FE ranged from 1 to 10 for the H-NS track, and from 1 to 1.8 for the Dps track. The GC content, calculated in a 25-bp sliding window, is shown as a heatmap.

**Supplementary Table S1.** Plasmids, strains, and primers. Lab collection IDs are assigned to all the materials.

| Plasmids | Description | Source |
|---|---|---|
| **pSC101 CmR plasmids for *in trans* expression of H-NS and Sfx from native promoters** | | |
| p450 | Empty pHSG576 | (15) |
| p451 | pHSG576-P*sfx*-*sfx*-FLAG | This work |
| p452 | pHSG576-P*sfx*-*sfx* | (16) |
| p487 | pHSG576-P*hns*-hns; pSC101 $Cm^R$ | (16) |
| **pRSF KanR plasmids for protein overexpression; T7 promoter-$His_{10}$-SUMO** | | |
| pIA1480 | Wild type H-NS overexpression vector | This work |
| pIA1481 | Wild type Hha overexpression vector | This work |
| pIA1649 | Wild type Sfx overexpression vector | This work |
| pIA1701 | Sfx with a unique C-terminal Cys | This work |
| pIA1720 | H-NS with a unique C-terminal Cys | This work |
| **R6K derivative plasmids; AmpR** | | |
| p103 | R6K wild type | (16) |
| pIA1548 | R6KΔ*sfx* | This work |
| pIA1682 | R6K75; the transposon region is removed except for the *bla* gene | This work |
| pIA1698 | R6K75Δ*sfx* | This work |
| pIA1710 | R6KΔ*sfx* with *yfjW* gene inserted into the ARG domain | This work |
| pIA1721 | R6K75*sfx*-FLAG; FLAG tag is added to the C-terminus of Sfx | This work |
| pIA1767 | P*actX*-*yfp* inserted in R6KΔ*sfx* | This work |
| pIA1768 | P*actX*-*yfp* inserted in R6K | This work |
| **Miscellaneous plasmids** | | |
| p28 | pBAD33 P15A $Cm^R$ | (17) |
| pDM-N1-400 | 5848 bp pBR322 backbone vector (Addgene #170474) | (18) |
| pIA1723 | P*actX*-*yfp* reporter; P*actX* cloned into pYPet-His (Addgene #14031) | This work |
| pIA1756 | *actX-sltX1-tivB2* fragment cloned into a pRSF plasmid | This work |
| **Strains** | **Genotype** | **Source** |
| IA294 | *Escherichia coli* K-12 BW25113 Keio collection wild type | (19) |
| IA357 | *E. coli* BW25113 Δ*hha*::$Kan^R$ | (19) |
| IA428 | *E. coli* BW25113 *lysA*::$Kan^R$ | (19) |
| IA765 | *Escherichia coli* K-12 MG1655 | Lydia Freddolino |
| IA969 | *E. coli* MG1655 *hns*-3xFLAG | Joseph Wade |
| IA1016 | *E. coli* MG1655 *F' lacZY*::Tn9 $Cm^R$ | Natacha Ruiz |
| IA1017 | IA1016 *rfaH*::$Kan^R$ | This work |
| IA1018 | IA1016 *hns*::$Kan^R$ | This work |
| IA1030 | *E. coli* BW25113 *argE*::$Tet^R$ | This work |
| BW55 | *E. coli* MG1655 *yfjw*::*actX-sltX1::*$Kan^R$ | This work |
| BW56 | *E. coli* MG1655 *hns*-3xFLAG *yfjw*::*actX-sltX1*::$Kan^R$ | This work |

| Primers | Sequences (5' – 3') | Usage |
|---|---|---|
| 3044 | CTTGATCGTTGGGAACCGGAG | *bla* qPCR |
| 3045 | TATCCGCCTCCATCCAGTCT | |
| 3063 | TCCTCCTTCTGGCTATCGGT | *tivB10* qPCR |
| 3064 | TGTCATCCCTCTGGCTCTGA | |
| 3069 | CGAAAGGAGGGCGATAGTGG | *sltX1* qPCR |
| 3070 | TTGTGCATGGCGAGAAAAGC | |
| 3086 | CCGGGGGTATTGTTGCTCTT | *tivB6* qPCR |
| 3087 | AATGCCAACGCACCGAAAAG | |
| 3079 | CTGAAGCAGCACGCAAAGAG | *Rho* qPCR |
| 3080 | GCGCAAAATAGCGTTCACCT | |
| 3113 | CCACGACAAGATTTGCAGCC | *actX* qPCR |
| 3114 | GAAGCCCGGTGATAGTGGTC | |
| 3325 | TCAACAGCATCTACATGATGGCC | *rpoC* qPCR |
| 3326 | CGCCATCAGACCACGCATAC | |
| 3335 | GCAAGCGATGGGGTGATTTT | *pir* qPCR |
| 3336 | ATATGGCGCTTGCTCCCATT | |
| 3343 | GGCGTGTTACGGTGAAAACC | *Cat* qPCR |
| 3344 | AAACTCACCCAGGGATTGGC | |
| 3490 | GGAGCATATGGCCTCGACTC | *ihfB* qPCR |
| 3491 | TCGCCAGTCTTCGGATTACG | |
| 3512 | TGGAGCAGGCCAATGAGAAG | *traV* qPCR |
| 3513 | GCATTGTCCGGAAGTTCCCT | |
| 3516 | AGGGAACGGCGAAAAAGGAT | *traU* qPCR |
| 3517 | TGTTCAGCCAGTACGTCAGC | |
| 3518 | CCGGGCTTATCATCATGGGG | *traS* qPCR |
| 3519 | CCATCCTCCGGCTAACACAG | |
| 3520 | AGGGCAGTGTCGATAAGGATG | *traI* qPCR |
| 3521 | GAGAAGGTCAGATCGTAGCCG | |
| 3552 | CAGACACACCAACGTATCTGGA | *yfjW* qPCR |
| 3553 | GGAAGGGCATAACAACCGGA | |
| 3554 | GGATAGTTAGTGGCGTTGCG | *waaL* qPCR |
| 3555 | GGGAACAGGAGTAGGGTTGC | |
| 2282 | GCTAGCACAAGGGAGTAGTTG | *actX-sltX1* EMSA fragment |
| 2771 | AGCAAGATTTGTGCATGGCG | |
| 2659 | CTAAAACAGTCGAAGTAACACC | PCR *vir* EMSA fragment |
| 3060 | CCCCGGCATTTTTCCCCATA | |
| 3465 | CAGTAATACGACTCACTATAGGGGCTGATTCTGGATTCGA | T7 promoter-*ssrA* template for making SsrA RNA |
| 3466 | TTTGGTGGAGCTGGCGGGAG | |

### References


1. Lu, J., Peng, Y., Wan, S., Frost, L.S., Raivio, T. and Glover, J.N.M. (2019) Cooperative Function of TraJ and ArcA in Regulating the F Plasmid tra Operon. *J Bacteriol*, **201**.
2. Will, W.R. and Frost, L.S. (2006) Characterization of the opposing roles of H-NS and TraJ in transcriptional regulation of the F-plasmid tra operon. *J Bacteriol*, **188**, 507-514.
3. Koraimann, G. (2018) Spread and Persistence of Virulence and Antibiotic Resistance Genes: A Ride on the F Plasmid Conjugation Module. *EcoSal Plus*, **8**.
4. Sanderson, K.E. and Stocker, B.A. (1981) Gene rfaH, which affects lipopolysaccharide core structure in Salmonella typhimurium, is required also for expression of F-factor functions. *J Bacteriol*, **146**, 535-541.
5. Wang, B. and Artsimovitch, I. (2020) NusG, an Ancient Yet Rapidly Evolving Transcription Factor. *Front Microbiol*, **11**, 619618.
6. Hustmyer, C.M., Wolfe, M.B., Welch, R.A. and Landick, R. (2022) RfaH Counter-Silences Inhibition of Transcript Elongation by H-NS-StpA Nucleoprotein Filaments in Pathogenic Escherichia coli. *mBio*, **13**, e0266222.
7. Wang, B., Mittermeier, M. and Artsimovitch, I. (2022) RfaH May Oppose Silencing by H-NS and YmoA Proteins during Transcription Elongation. *J Bacteriol*, **204**, e0059921.
8. Sevostyanova, A., Belogurov, G.A., Mooney, R.A., Landick, R. and Artsimovitch, I. (2011) The beta subunit gate loop is required for RNA polymerase modification by RfaH and NusG. *Mol Cell*, **43**, 253-262.
9. Lartigue, M.F., Leflon-Guibout, V., Poirel, L., Nordmann, P. and Nicolas-Chanoine, M.H. (2002) Promoters P3, Pa/Pb, P4, and P5 upstream from bla(TEM) genes and their relationship to beta-lactam resistance. *Antimicrob Agents Chemother*, **46**, 4035-4037.
10. Reese, M.G. (2001) Application of a time-delay neural network to promoter annotation in the Drosophila melanogaster genome. *Comput Chem*, **26**, 51-56.
11. Shen, B.A., Hustmyer, C.M., Roston, D., Wolfe, M.B. and Landick, R. (2022) Bacterial H-NS contacts DNA at the same irregularly spaced sites in both bridged and hemi-sequestered linear filaments. *iScience*, **25**.
12. Gavrilov, A.A., Shamovsky, I., Zhegalova, I., Proshkin, S., Shamovsky, Y., Evko, G., Epshtein, V., Rasouly, A., Blavatnik, A., Lahiri, S. *et al.* (2025) Elementary 3D organization of active and silenced E. coli genome. *Nature*, **645**, 1060-1070.
13. Ali, S.S., Whitney, J.C., Stevenson, J., Robinson, H., Howell, P.L. and Navarre, W.W. (2013) Structural insights into the regulation of foreign genes in Salmonella by the Hha/H-NS complex. *J Biol Chem*, **288**, 13356-13369.
14. McCarthy, L.A., Way, L.E., Dai, X., Ren, Z., Fuller, D.E.H., Dhiman, I., Larkin, L., Sieben, J.J.D., Westerlaken, I., Abbondanzieri, E.A. *et al.* (2026) Dps binds and protects DNA in starved Escherichia coli with minimal effect on chromosome accessibility, dynamics, and organization. *Nucleic Acids Research*, **54**.
15. Takeshita, S., Sato, M., Toba, M., Masahashi, W. and Hashimoto-Gotoh, T. (1987) High-copy-number and low-copy-number plasmid vectors for lacZ alpha-complementation and chloramphenicol- or kanamycin-resistance selection. *Gene*, **61**, 63-74.
16. Wang, A., Cordova, M. and Navarre, W.W. (2025) Evolutionary and functional divergence of Sfx, a plasmid-encoded H-NS homolog, underlies the regulation of IncX plasmid conjugation. *mBio*, **16**, e02089-02024.
17. Guzman, L.M., Belin, D., Carson, M.J. and Beckwith, J. (1995) Tight regulation, modulation, and high-level expression by vectors containing the arabinose PBAD promoter. *J Bacteriol*, **177**, 4121-4130.
18. Qian, J., Wang, B., Artsimovitch, I., Dunlap, D. and Finzi, L. (2024) Force and the α-C-terminal domains bias RNA polymerase recycling. *Nat Commun*, **15**, 7520.

19. Baba, T., Ara, T., Hasegawa, M., Takai, Y., Okumura, Y., Baba, M., Datsenko, K.A., Tomita, M., Wanner, B.L. and Mori, H. (2006) Construction of Escherichia coli K-12 in-frame, single-gene knockout mutants: the Keio collection. *Mol Syst Biol*, **2**, 2006.0008.
